# Light quality affects chlorophyll biosynthesis and photosynthetic performance in Antarctic Chlamydomonas

**DOI:** 10.1101/2024.08.01.606229

**Authors:** Mackenzie C. Poirier, Kassandra Fugard, Marina Cvetkovska

## Abstract

The perennially ice-covered Lake Bonney in Antarctica has been deemed a natural laboratory for studying life at the extreme. Photosynthetic algae dominate the lake food webs and are adapted to a multitude of extreme conditions including perpetual shading even at the height of the austral summer. Here we examine how the unique light environment in Lake Bonney influences the physiology of two *Chlamydomonas* species. *Chlamydomonas priscuii* is found exclusively in the deep photic zone where is receives very low light levels biased in the blue part of the spectrum (400-500 nm). In contrast, *Chlamydomonas* sp. ICE-MDV is represented at various depths within the water column (including the bright surface waters), and it receives a broad range of light levels and spectral wavelengths. The close phylogenetic relationship and psychrophilic character of both species makes them an ideal system to study the effects of light quality and quantity on chlorophyll biosynthesis and photosynthetic performance in extreme conditions. We show that the shade-adapted *C. priscuii* exhibits a decreased ability to accumulate chlorophyll and severe photoinhibition when grown under red light compared to blue light. These effects are particularly pronounced under red light of higher intensity, suggesting a loss of capability to acclimate to varied light conditions. In contrast, ICE-MDV has retained the ability to synthesize chlorophyll and maintain photosynthetic efficiency under a broader range of light conditions. Our findings provide insights into the mechanisms of photosynthesis under extreme conditions, and have implications on algal survival in changing conditions of Antarctic ice-covered lakes.

## Introduction

Light is an essential energy source for the fixation of atmospheric CO_2_ into organic carbon forms, but photosynthetic organisms are faced with variable and often rapidly changing light environments. The capture of light by chlorophyll (Chl) in the light harvesting antenna complexes (LHCs) is the first step in this process of biological carbon fixation. Green algae have two types of chlorophyll (Chl *a* and Chl *b*), with maximal absorbance in the blue (∼400-500 nm) and red (∼600-700 nm) part of the visible spectrum. Chl *a* is found in the reaction centers of photosystem I (PSI) and photosystem II (PSII), as well as within the LHCs associated with each photosystem while Chl *b* is only present in the antennae. Photosynthetic complexes require strict Chl *a/b* stoichiometry for optimal energy transfer at different light conditions (Tanaka and Tanaka 2007). Furthermore, the presence of Chl *b* is necessary for the correct assembly, accumulation, and stability of LHCs in algae (Bujaldon et al. 2017), bryophytes (Zhang et al. 2023) and angiosperms (Kim et al. 2009; Krόl et al. 1995; Nick et al. 2013; Reinbothe et al. 2006;). Thus, the ability to accumulate stable levels of Chl and appropriate Chl *a/b* ratios are key requirements for acclimating to variable light.

Chl is synthesized in a complex multi-step pathway, tightly regulated by the availability and spectral composition of light (reviewed by Kobayashi and Masuda 2019; Masuda and Fujita 2008; Yong et al. 2024). The penultimate regulatory step in this pathway is the reduction of protochlorophyllide (Pchlide) to chlrophyllide *a* (Chlide *a*), a direct precursor to Chl *a*. This reaction is catalyzed by two nonhomologous enzymes: light-dependent (LPOR) and dark-operative (DPOR) protochlorophyllide oxidoreductases (Masuda and Yuichi 2008; Vedelankar and Tripathy 2019). The nuclear-encoded LPOR is ubiquitously present in all photosynthetic eukaryotes and is only active when Pchlide absorbs light (Griffiths et al. 1996; Shui et al. 2009). The plastid-encoded protein complex DPOR, on the other hand, has been lost multiple times independently throughout eukaryotic evolution and does not require light for activity (Hunsperger et al. 2015; Kim et al. 2017; Shui et al. 2009). Chl *b* is derived from Chl *a* solely through the action of the enzyme chlorophyllide *a* oxygenase (CAO; Tanaka et al. 1998). The details of CAO regulation are not fully understood, but it is known that light intensity regulates CAO gene expression (Biswal et al. 2012; Biswal et al. 2023; Pattanayak et al. 2005; Tanaka and Tanaka 2005) and the accumulation of its product Chl *b* regulates CAO protein stability (Nakagawara et al. 2007; Sakuraba et al. 2009; Yamasato et al. 2005). These crucial steps influence the total availability of chlorophylls, Chl *a/b* ratios, and consequently, the size of the LHCs.

Many of the intricacies of how light intensity and spectral wavelengths affect green algae come from studies on the model *Chlamydomonas reinhardtii*, an alga well adapted to a wide range of light conditions (Li et al. 2023; Salomé et al. 2019). Green algae can employ multiple strategies to optimize light absorption and minimize light-induced damage to the photosynthetic apparatus when exposed to variable light conditions. Short-term acclimation occurs on the time scale of seconds to minutes and includes the dissipation of excess light energy as heat through non-photochemical quenching (NPQ), balancing the excitation state of PSII and PSI through state transitions, inducing alternative electron routes through cyclic electron flow (CEF), and phototactic movement towards or away from light. Long-term acclimation happens on the time scale of hours to days through changes in gene expression and protein accumulation of key photosynthetic proteins that ultimately alter the composition of pigments, accumulation of LHCs, and the size of the photosynthetic complexes (reviewed in Allorent and Petroutsos 2017; Erickson et al. 2015; Lu et al. 2022; Pinnola 2019). *C. reinhardtii* has been crucial for elucidating many photosynthetic traits but this temperate species lacks the capacity for survival under extreme conditions (Sasso et al. 2018). Algae are ubiquitous and found in many environments that are unsuitable for the growth of most models (Rappaport and Oliverio 2023), and the mechanisms behind light perception and energy balance in these non-model species are yet to be understood.

More than 70% of Earth’s biosphere is permanently cold (<5°C). Virtually all cold-water food webs are supported by photosynthetic microbes, including eukaryotic alga (De Maayer et al. 2014). Lake Bonney is part of the McMurdo Long Term Ecological Research in Antarctica and has been studied as a natural laboratory for life at the extreme since 1993 (mcmlter.org). The perennial ice-cover prevents wind-driven mixing and environmental input resulting in a very stable environment. The microbial food webs in the lake are dominated by vertically stratified and diverse photosynthetic communities (Bielewicz et al. 2011; Li and Morgan-Kiss 2019) adapted to life at perpetually low temperatures, nutrient deficiencies, high salinity and high oxygenation levels. In addition, photosynthetic algae in the lake are challenged with extreme shading under the ice even at the peak of the austral day, narrow spectral range (450-550 nm) due to the attenuation of higher wavelengths by the water column, and perpetual darkness during the austral night. These psychrophiles (cold extremophiles) truly push the boundaries of photosynthetic life.

Two psychrophilic Chlamydomonadalean species have recently emerged as excellent systems to study algal adaptation to extreme conditions. *Chlamydomonas priscuii* (previously UWO241) has been studied for more than 30 years (reviewed in detail in Cvetkovska et al. 2022a; Dolhi et al. 2013; Hüner et al. 2022) and was propelled to the status of model psychrophile by the recent sequencing of its genome (Zhang et al. 2021). This alga is found exclusively in the deep photic zone (17-18m) of Lake Bonney (Neale and Priscu 1995), where is receives very low light levels (∼10 µmol photons m^-2^s^-1^) biased in the blue-green spectrum (400-500 nm). Chlamydomonas sp. ICE-MDV is the dominant chlorophyte in the upper layers of the lake and has been detected at various depths (5-15 m; Li and Morgan-Kiss 2019) including shallow seasonally open-water moats between the two lobes of the lake (Sherwell et al. 2022). Thus, ICE-MDV is represented at a broad range of light levels and spectral wavelengths, including the high light ice-free surface waters of Lake Bonney (∼190 µmol m^-2^ s^-1^) (Sherwell et al. 2022).

Life at the extreme has influenced the physiology of both species, but to a different extent. *C. priscuii* has a ‘locked’ physiology and largely lacks the capacity to respond to short-term environmental challenges. This alga has an attenuated heat stress response at non-permissive temperatures (Cvetkovska et al. 2022b), minimal PSI restructuring under iron stress (Cook et al. 2019), has lost the ability to balance PSI and PSII light excitation through state transitions (Morgan-Kiss et al. 2002; Szyszka-Mroz et al. 2015) and has a very weak phototactic response (Poirier et al. 2023). Curiously, *C. priscuii* lacks the genes that encode for DPOR and thus relies solely on LPOR to synthesize chlorophyll (Cvetkovska et al. 2018a). In contrast, ICE-MDV displays a more flexible physiology and a photosynthetic apparatus similar to its non-extremophilic relatives. ICE-MDV has retained the capacity for state transitions, PSI restructuring and phototaxis (Cook et al. 2019; Kalra et al. 2023; Poirier et al. 2023), and can induce protective mechanisms under temperature stress (Cvetkovska et al. 2022b). Its plastid genome does encode for DPOR, giving this alga the flexibility of both light dependent and independent chlorophyll biosynthesis (Smith et al. 2019). The reason behind these different physiologies is not known, but it has been suggested that ICE-MDV is a more recent arrival in Lake Bonney that has not yet been constrained by the highly stable conditions underneath the ice (Smith et al. 2019).

Lake Bonney is one of the last perennially ice-covered and pristine lakes on our planet, but even this isolated environment has suffered anthropogenic disturbances. The permanent ice cover has thinned in the past decades leading to the appearance of open water perimeter moats (Gooseff et al. 2017; Obryk et al. 2019), which has had a profound effect on the shade-adapted phytoplankton communities. Field and experimental studies on native Lake Bonney microbiota demonstrated that shade-adapted phytoplankton were severely impacted by the higher light conditions present in open-water moats. The increased light intensity caused inhibition of photochemistry and downregulation of the photosynthetic apparatus, leading to population declines, particularly evident in the Chlorophyta (Sherwell et al. 2022).

Changing ice cover alters not only the intensity but also the spectral wavelengths of available light (Howard-Williams et al. 1998). Building on these insights, we tested the impact of light intensity and spectral wavelengths on two psychrophilic species native to the lake. We hypothesized that the dominant lake Chlorophyte ICE-MDV found throughout the water column (including seasonally open waters), will be better equipped to acclimate to light variability when compared with the shade-adapted ‘denizen of the deep’ *C. priscuii*.

## Materials and Methods

### Growth conditions

*Chlamydomonas priscuii* (previously UWO241; strain CCMP1619) and *Chlamydomonas sp.* ICE-MDV were grown axenically in Bold’s Basal Medium (BBM) supplemented with 70 mM NaCl at 8°C, conditions previously shown to support robust growth and high photosynthetic efficiency (Szyszka et al. 2007). Cultures were grown under continuous light provided by LED light bulbs. Light intensity was either ∼100 µmol photons m^-2^ s^-1^ (increased light, standard lab culturing conditions; Hui et al. 2023) or ∼10 µmol photons m^-2^ s^-1^ (low light, equivalent to light levels at ∼15-m depth in Lake Bonney; Lizotte and Priscu 1992). Light intensity was measured with a quantum sensor attached to a radiometer (Model LI-189; Li-COR). Light quality was determined by spectroscopic measurements (Fig. 1; HR2 VIS-NIR Spectrophotometer, OceanOptics). Blue light (420-500 nm) was chosen to resemble the light quality in the under-ice aquatic habitats of Lake Bonney. Conversely, red light (620-690) is limited in these environments and represents a light quality that these species do not naturally experience (Neale and Priscu 1995). Cultures were grown in 250 mL glass Pyrex tubes in a temperature regulated aquaria and aerated with sterile ambient air provided by aquarium pumps. Experimental cultures were seeded and acclimated to the respective light conditions prior to measurements. All experiments were done on actively growing cultures during mid-log phase of exponential growth.

**Fig. 1.**
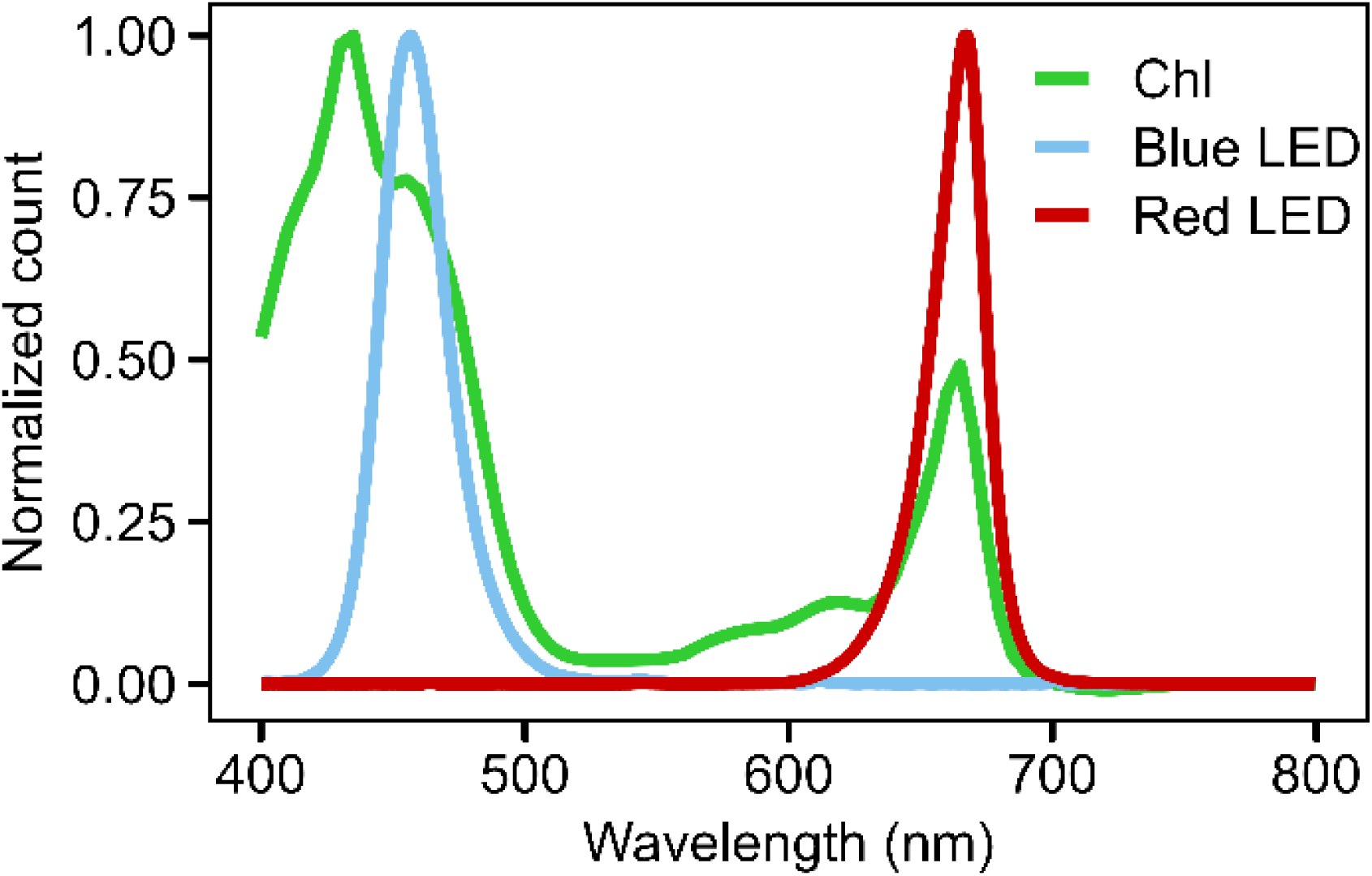
The spectral distribution of LED light sources used in this study. Absorption spectra of algal chlorophyll extracted in 80% acetone is also provided to demonstrate the overlap with the major light peaks examined in this study (Blue: 420-500 nm; Red: 620-690 nm). Normalized count values are a measure of signal intensity produced by the light source at a given wavelength

### Growth measurements and chlorophyll quantification

Algal growth was monitored by optical density (OD), chlorophyll content, and cell counts. Change in OD was measured spectrophotometrically at 750 nm (Cary 60 UV-Vis spectrophotometer, Agilent Technologies). Chl *a* and *b* concentrations were determined from whole cell extracts in 90% acetone and measured spectrophotometrically at 647 and 665 nm. Pigment content was calculated according to Jeffery and Humphrey (1975). Cell numbers and their approximate size were determined with a Countess II FL Automatic Cell Counter (ThermoFisher Scientific) using brightfield imaging. Cell death was measured with the fluorescent dye SYTOX Green (Ex/Em = 504/523 nm; ThermoFisher Scientific) that accumulates in dead cells only. In all cases, 2 µM SYTOX Green was added to culture samples, followed by 15 min dark incubation prior to visualization. Cell death was quantified as a proportion of green-fluorescing cells by fluorescence imaging (EVOS™ Cy5 and GFP Light Cubes, ThermoFisher Scientific). Growth curves were produced by graphing OD_750_ as a function of time. Growth rate was calculated using the natural log transformation of OD_750_ values during exponential growth according to the following formula (t = time; x = OD_750_ at that given time point):

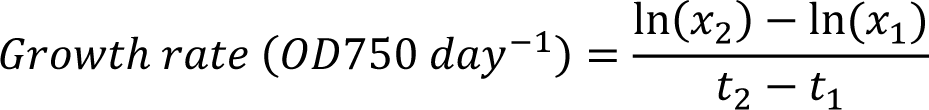

### Gene identification

The *C. priscuii* nuclear and plastid genomes were recently sequenced (Zhang et al. 2021; Cvetkovska et al. 2018a). The genome of ICE-MDV has been sequenced (Raymond and Morgan-Kiss 2017) but is not fully assembled and annotated (331,087 contigs). Thus, we used the ICE-MDV data in combination with the published genome of the closely related Antarctic sea-ice alga *Chlamydomonas* sp. ICE-L (Zhang et al. 2020) to identify full-length homologs of our genes of interest. ICE-MDV and ICE-L share ∼100% identity in the sequences of several genes (Cook et al. 2019; Smith et al. 2019), indicating that these algae are very closely related or represent different strains of the same species. These datasets were screened for the presence of chlorophyll biosynthesis and light harvesting genes.

The full-length sequences for the genes encoding LPOR and CAO in the genome of *C. priscuii* and DPOR in the genome of ICE-MDV were previously identified (Cvetkovska et al. 2018a; Smith et al. 2019). In all other cases, we used sequences from *C. reinhardtii* (Craig et al. 2023; Merchant et al. 2007; Table S1) and conserved key domains obtained from Phytozome (Goodstein et al. 2012) as queries. Gene sequences were identified through tBLASTn searches (e-value < e^-10^) and manually inspected for redundancy and accuracy. The genomes of closely related algae were obtained from PhycoCosm (Grigoriev et al. 2021). Multiple sequence alignments were done with Clustal Omega (Sievers et al. 2011). Phylogenetic trees were constructed in MEGA XI (Kumar et al. 2018), with bootstrap confidence values calculated from 1000 iterations. Visualization was done in iTOL (v6.8.2; Letunic and Bork 2024).

### Gene expression

Gene specific primers (Table S1) were developed with Primer3 (v2.3.7; Untergasser et al. 2012). Since the genes encoding for the three DPOR subunits were not detected in the *C. priscuii* plastid genome (Cvetkovska et al. 2018a), we designed primers to highly conserved regions of these genes within several related species (ICE-MDV, *C. reinhardtii*, *C. eustigma*, *V. carteri*). To obtain RNA, algal cells were collected by centrifugation (5 000 g, 5 minutes), flash frozen in liquid nitrogen and stored at -80°C. RNA was extracted from frozen algal pellets using a modified CTAB protocol (Possmayer et al. 2016) and genomic DNA was removed using the Ambion TURBO DNA-free kit (ThermoFisher Scientific). The concentration and quality of RNA was determined spectrophotometrically using a Nanodrop 2000 (ThermoFisher Scientific). cDNA was synthesized with the GoScript Reverse Transcriptase kit following the manufacturer’s instructions (Promega). Droplet digital PCR (QX200 ddPCR; Bio-Rad) with EvaGreen Supermix (Bio-Rad) was used to quantify gene expression. Droplet generation was done according to manufacturer’s instructions (QX200 Droplet Generator; Bio-Rad). PCR cycling conditions were as follows: 1) 1x 95°C for 5 minutes; 2) 50x of a three-step cycling protocol (95°C for 30 seconds, 58°C for 1 minute, and 72°C for 30 seconds); 3) 1x 90°C for 5 minutes; 4) 1x 4°C for 5 minutes. Droplet reading was done using absolute quantification and results were analyzed with QuantaSoft^TM^ Software (Bio-Rad). Gene expression values are represented as gene copies µL^-1^ in the original cDNA sample. All experiments were performed with three biological replicates.

### SDS-PAGE and immunoblotting

Protein quantification was carried out based on the methods by Cook et al. 2019 and Szyszka-Mroz et al. 2015. In brief, total proteins were extracted from frozen cells by resuspending the pellet in 1.1 M Na_2_CO_3_ then freezing at -20°C for 1 hour. Protein samples were solubilized by adding an equal volume of Solubilization buffer (5% (w/v) SDS, 30% (w/v) sucrose) and heated for 5 minutes at 85°C. Protein concentration was quantified with a BCA protein Assay Kit according to manufacturer’s instructions (Bio Basics). Proteins were loaded on an equal protein basis (10 µg), separated on a 12.5% (v/v) SDS-PAGE gel as described previously (Szyszka-Mroz et al. 2015), and transferred to 0.2 µm PVDF membrane (Bio-Rad) using the Trans-Blot Turbo Transfer System (Bio-Rad). Transferred membranes were blocked with TBS-T containing 5% (w/v) non-fat milk overnight and probed with primary antibodies (Agrisera, Vännäs, Sweden) raised against proteins from the model alga *C. reinhardtii*: Lhcbm5 (1:5000, product #AS09 408), PsbA-D1 (1:15 000, product #AS05 084A), and PsaA (1:10 000, product #AS06 172). These antibodies have been confirmed to bind to their *C. priscuii* and ICE-MDV homologs previously (Cook et al. 2019). Membranes were then exposed to HRP-conjugated secondary antibody for 1 hour (1:10 000 Goat anti-rabbit IgG HRP conjugate; Bio-Rad). Antibody-protein complexes were visualized using enhanced chemiluminescence detection reagents (Clarity Western ECL Substrate, Bio-Rad) and imaged using the ChemiDoc^TM^ Imaging System (Bio-Rad). All experiments were done in at least three biological replicates.

### Room temperature chlorophyll *a* fluorescence

*In vivo* room temperature Chl *a* fluorescence was measured using a pulse amplitude modulated chlorophyll fluorometer with a DR detector (DUAL-PAM-100; Walz, Germany). Live cultures (1 mL; 3.5 µg Chl/mL) were stirred with a magnetic stir bar and kept at constant temperature (8°C) using a water-jacketed quartz cuvette connected to an external water bath. After 10 minutes of dark adaptation, Chl fluorescence at open PSII reaction centers (F_o_) was measured by excitation with a non-actinic measuring beam (0.13 µE) pulsed at 20 Hz. Maximum fluorescence at closed PSII reaction centers (F_m_) was induced by a saturating pulse (200 ms, 5325 μE). A rapid light curve was produced using actinic light increasing from 6 µE to 1763 µE based on methods by Ralph and Gademann (2005). The period for each light intensity was 10 seconds with saturating light pulses (200 ms, 10 505 μE) after each step. The same cells were used to calculate dark relaxation kinetics by turning off the light and giving a saturation pulse (200 ms, 10 505 µE) after 30 seconds, 1 minute, 2 minutes, 5 minutes, and 10 minutes in the dark. Data acquisition was managed using the WinControl software (Walz, Germany). All experiments were performed as at least three biological replicates. Fluorescence parameters were calculated using the equations described by Kramer et al. (2004) and Kitajima and Butler (1975). Maximum PSII photochemical efficiency (F_V_/F_M_) was calculated as F_M_–F_o_/F_M_, using dark-adapted cells (Kitajima and Butler 1975). The quantum yield of steady-state photosynthesis was calculated as: Y(II) = (F_M_′–F_S_)/F_M_′, Y(NPQ) = (F/F_M_′)-(F/F_M_), Y(NO) = F/F_M_, where Y(II) is the yield of PSII photochemistry, Y(NPQ) is the yield of non-photochemical energy dissipation by down-regulation through antenna quenching, and Y(NO) is the yield of all other processes involved in non-photochemical energy losses (Kramer et al. 2004). The curve of relative electron transport rate (ETR) at PSII during the rapid light curve was fit using the Platt, Gallegos, and Harrison 1980 model (Platt et al. 1980) and the phytotools R package (Revell 2024) to determine the light harvesting efficiency (α), light-saturated electron transport rate (ETR_max_), and minimum saturating irradiance (Ek) values.

### Statistical analysis

All statistics were performed using R. For growth rates, Chl measurements, chlorophyll *a* fluorescence, and cell size data a multi-way ANOVA was performed with three independent variables of light quantity, light quality, and species. For gene expression data a two-way ANOVA was performed for each species with two independent variables of light quantity and light quality. Tukey’s HSD post-hoc test was used in each case to identify significant differences (p<0.05) between conditions.

## Results

### The effect of light spectral quality on growth physiology

Both *C. priscuii* and ICE-MDV originate from different depths within the water column of the ice-covered Lake Bonney (Lizotte and Priscui 1992). We demonstrated that the growth rate of these species in a closed culture was significantly dependent on both light quality and quantity (Fig. 2a). Both *C. priscuii* and ICE-MDV exhibited the highest maximal growth rates under higher intensity blue light (0.63 and 0.56 ΔOD_750_ d^-1^, respectively) when compared to lower intensity blue light (0.38 and 0.35 ΔOD_750_ d^-1^, respectively); however, we observed significantly different responses when these species were cultured under red light (Fig. 2a). While *C. priscuii* demonstrated only a small growth decrease in red light compared to blue light, we observed very low maximal growth rates in ICE-MDV in red light, regardless of the light intensity (0.13-0.20 ΔOD_750_ d^-1^; Fig. 2a). Furthermore, we demonstrate that in all cases cellular death was <10% (Fig. 2b), indicating that the low apparent growth is not due to algal death.

**Fig. 2.**
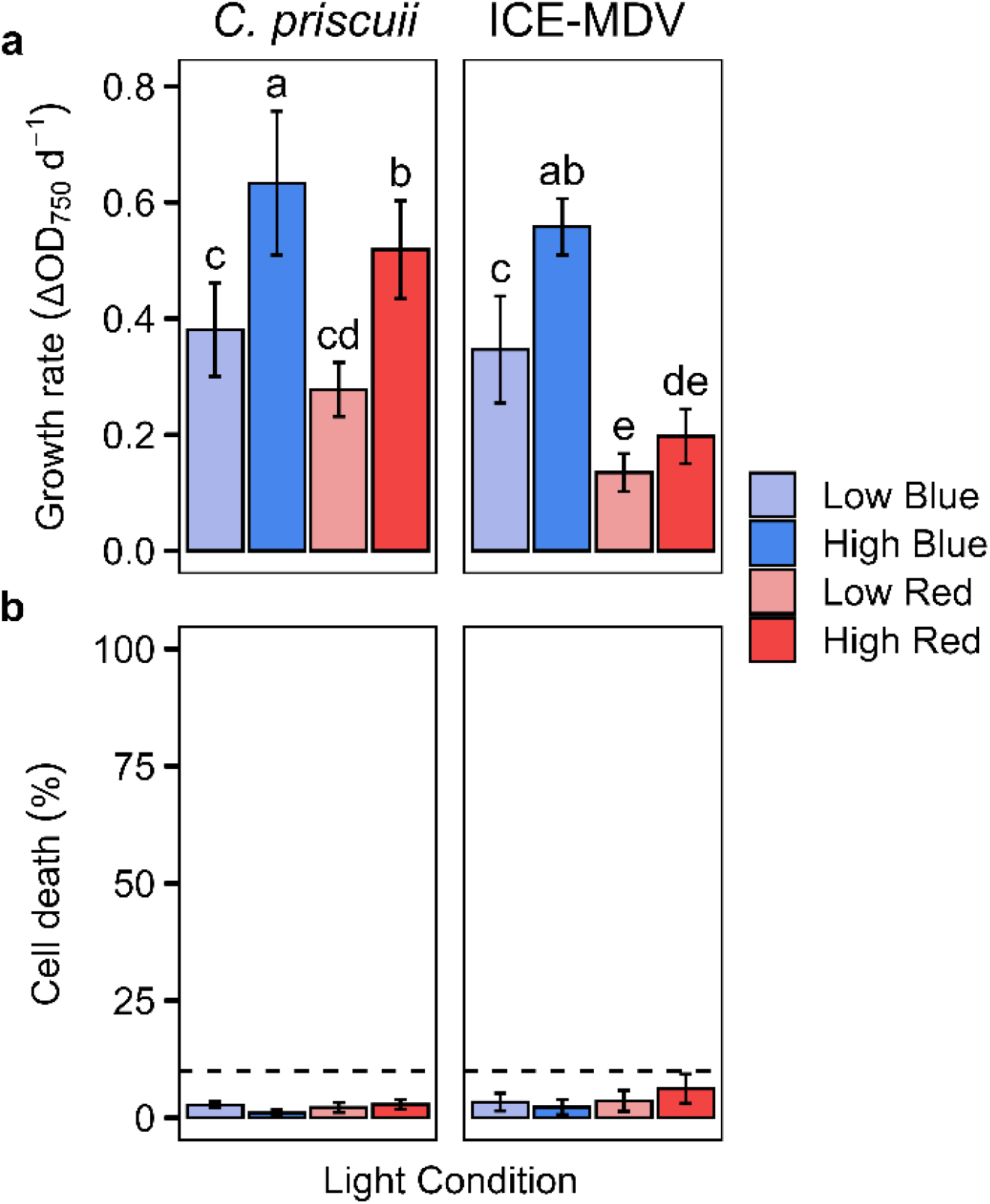
Maximal growth rates (a) and cell death (b) of exponentially growing *C. priscuii* and ICE-MDV cultures exposed to different steady state light conditions. Growth rates were determined based on changes in optical density (750 nm). Dashed line indicates 10% cell death. Statistically significant differences are represented as different letters as determined using Tukey’s post hoc test (p < 0.05). Values are means ± SD (n ≥ 9)

Next, we examined the ability of the algae to accumulate Chl and to adjust the Chl *a/b* ratios under different light conditions. In accordance with previous work (Stahl-Rommel et al. 2021; Szyszka et al. 2007), we show that both psychrophiles have more total Chl and lower Chl *a/b* ratios when grown under lower light intensity (Fig. 3). Acclimation to higher levels of blue light leads to a decrease in total Chl in both species (Fig. 3a), accompanied with higher Chl *a/b* ratios (Fig. 3b). The most striking difference between species was observed in cultures grown under red light. *C. priscuii* maintained the ability to adjust cellular Chl *a/b* ratios but exhibited very low total Chl levels, regardless of the light intensity (0.74-1.11 pg/cell; Fig. 3a). In contrast, ICE-MDV exhibited low Chl levels under higher intensity red light (1.22 pg/cell) but retained the ability to accumulate more total Chl in lower intensity red light (4.13 pg/cell).

**Fig. 3.**
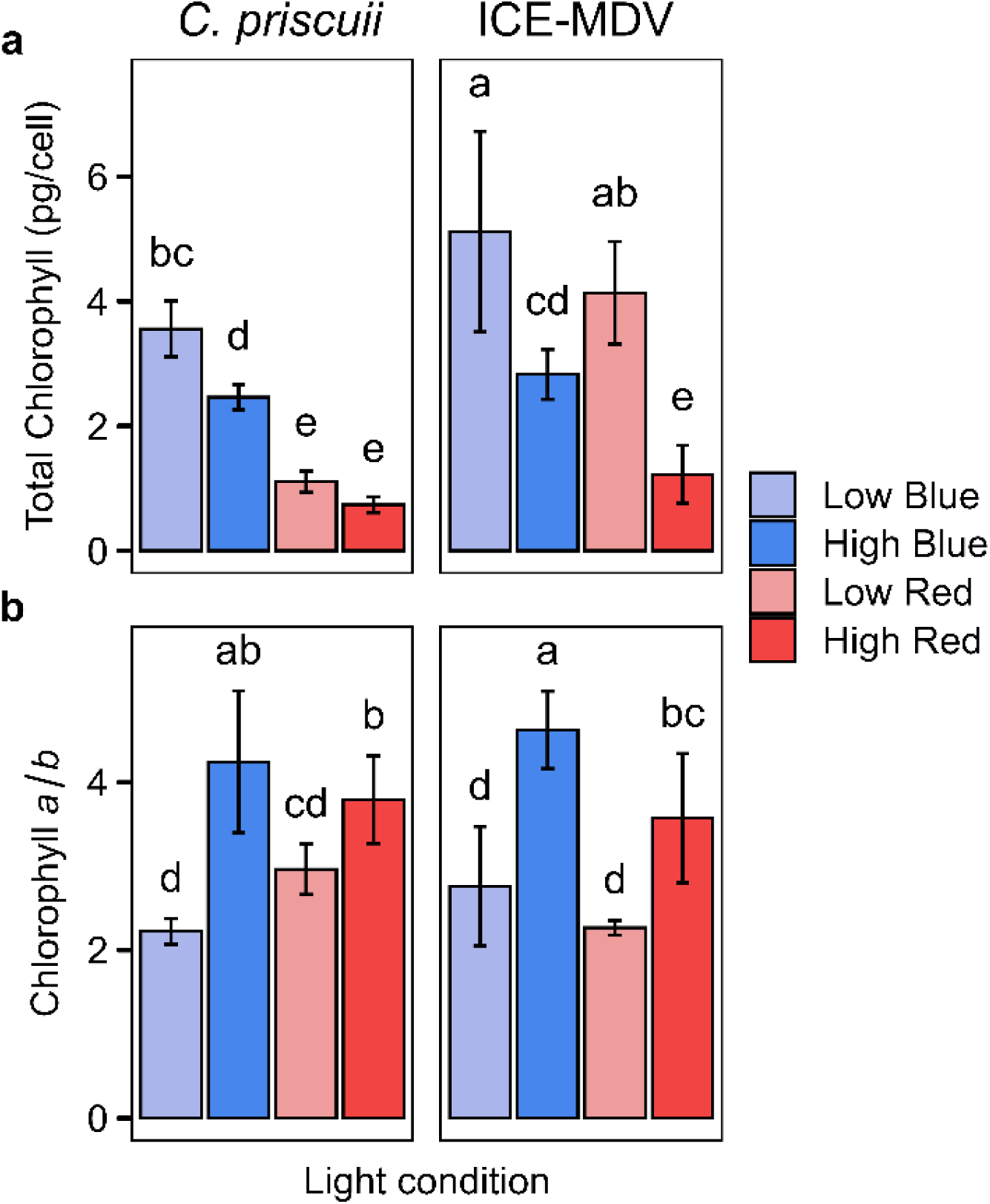
Chlorophyll accumulation in C. priscuii and ICE-MDV grown under different steady state light conditions. Chl per cell levels (a) and Chl a/b ratios (b) determined in exponentially growing C. priscuii and ICE-MDV cultures. Statistically significant differences are represented as different letters as determined using Tukey’s post hoc test (p < 0.05). Values are means ± SD (n ≥ 9)

The differences in cellular Chl levels were not due to changes in cell size. While *C. priscuii* cells are generally smaller (6.1-9.1 µm) compared to ICE-MDV (9.7-11.6 µm), we observed very little difference in cell size in cultures acclimated to different light conditions (Fig. S1).

### The presence and expression of key genes in the chlorophyll biosynthesis pathway

To gain better insight into the processes that underlie chlorophyll accumulation, we examined the key components of the Chl biosynthesis pathway at the level of gDNA and mRNA. Examination of the *C. priscuii* and ICE-MDV genomes revealed a conserved Chl biosynthesis pathway (Table S2) with a few notable exceptions. First, previous computational analyses suggested that *C. priscuii*, but not ICE-MDV, lacks the plastid-encoded genes encoding all three subunits of DPOR (*chlN, chlL,* and *chlB*) (Cvetkovska et al. 2018a; Smith et al. 2019). We experimentally confirmed the lack of DPOR in *C. priscuii* and its presence in ICE-MDV by PCR amplification using primers designed to bind highly conserved regions of *chlN* and *chlL* (Fig. 4). This analysis further supports the hypothesis that *C. priscuii* relies solely on the light-dependent activity of LPOR to synthesize Chl (Cvetkovska et al. 2018a). It should be noted that the *chlB* subunit was not used to test for the presence of DPOR as this subunit is less conserved among species and it was not possible to design conserved primers. We also identified the gene encoding for LPOR in ICE-MDV and related chlorophytes. We show that the gene encoding for LPOR in *C. priscuii* had the highest homology to the LPOR gene encoded in the ICE-MDV genome (77.5% identity) and the acidophile *Chlamydomonas eustigma* (81.4% identity), including a short 24 amino acid domain not observed in non-extremophilic chlorophytes (Fig. S2).

**Fig. 4.**
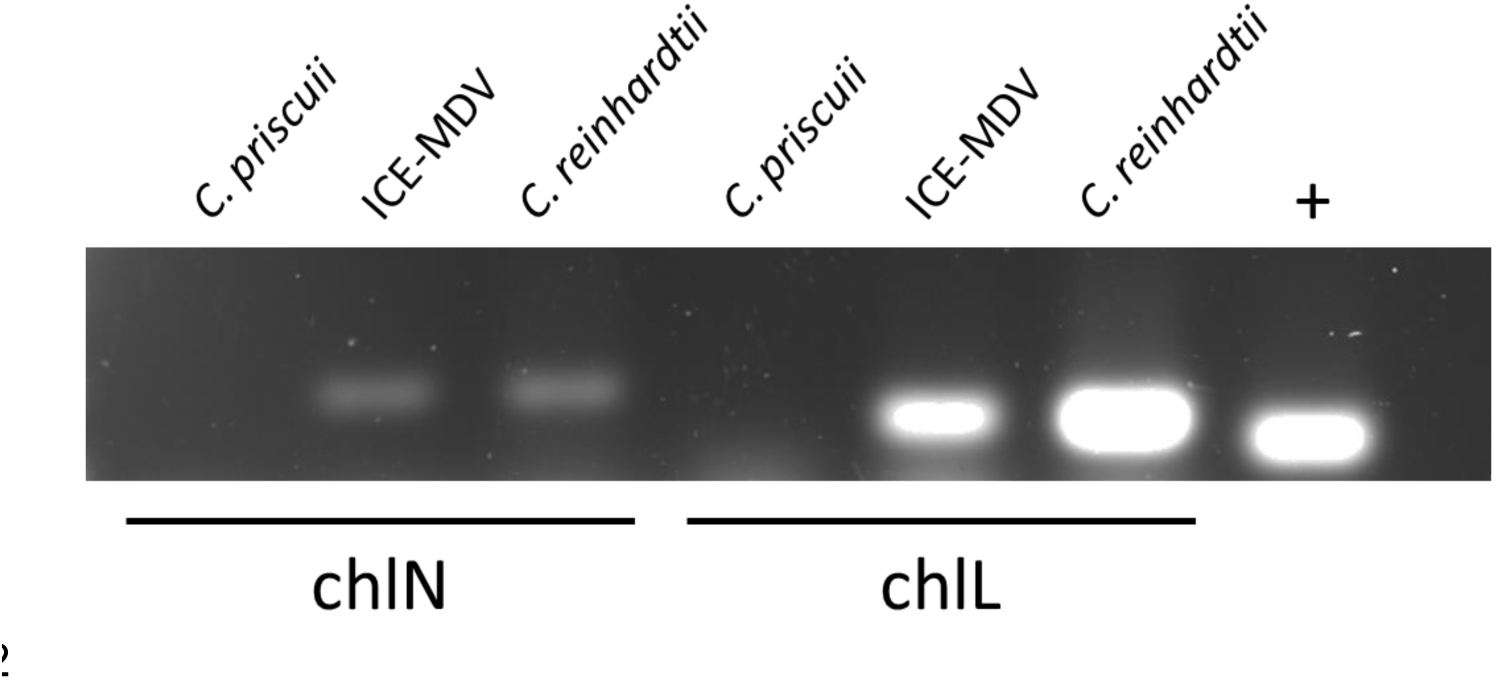
Experimental evidence for the absence of DPOR in *C. priscuii* and its presence in ICE-MDV. Primers designed to bind to conserved regions of the genes encoding for the DPOR subunits in several algal species were able to amplify regions in chlN and chlL in ICE-MDV and *C. reinhardtii* but not in *C. priscuii*. A PCR reaction using primers specific to CAO1 (a gene that is present in all three algal species) was used as a positive control (+) with *C. priscuii* cDNA to confirm absence of DPOR bands was not caused by technical errors

Detailed examination of the ICE-MDV genome revealed two copies of the gene that encodes for CAO (Fig. S3), the enzyme that catalyzes the conversion of Chl *a* to Chl *b* (Tanaka and Tanaka 2019). Significantly, the only other known green alga that also harbours a CAO duplication in its genome is *C. priscuii* (Cvetkovska et al. 2018). To confirm that the detected CAO duplicates were not an artifact in the genome assemblies, these genes were amplified from both species and sequenced, confirming the presence of two CAO homologs experimentally (not shown). *C. priscuii* and ICE-MDV CAO1 and CAO2 are more similar within the species (71.8% and 75.6% identity, respectively) than *C. priscuii* CAO genes are to ICE-MDV CAO genes (59.1% and 65% identity) suggesting that the duplication event occurred independently within each species (Fig. S3).

To understand the importance of these key genes in chlorophyll biosynthesis in different light conditions we examined their expression using ddPCR, a method that allows for absolute quantification of transcript copies. LPOR expression did not change significantly in any of the examined light conditions in either species (Fig. 5a). The expression of *chlN*, used as a proxy for DPOR expression in ICE-MDV, was also not significantly affected by light treatment (Fig. 5a). This suggests that Chl accumulation, while affected by light quality, is not regulated at the level of LPOR and DPOR expression. Our data also revealed another trend. LPOR expression was on average lower in ICE-MDV compared to *C. priscuii* (∼6 000 copies/µL vs ∼11 000 copies/µl, respectively). However, we show that the combined expression of DPOR and LPOR in ICE-MDV was on par with the expression of LPOR alone in *C. priscuii* (∼11,000 copies/μL; Fig. 5a). Thus, it is possible that *C. priscuii* compensates for the lack of DPOR by increasing LPOR expression.

**Fig. 5.**
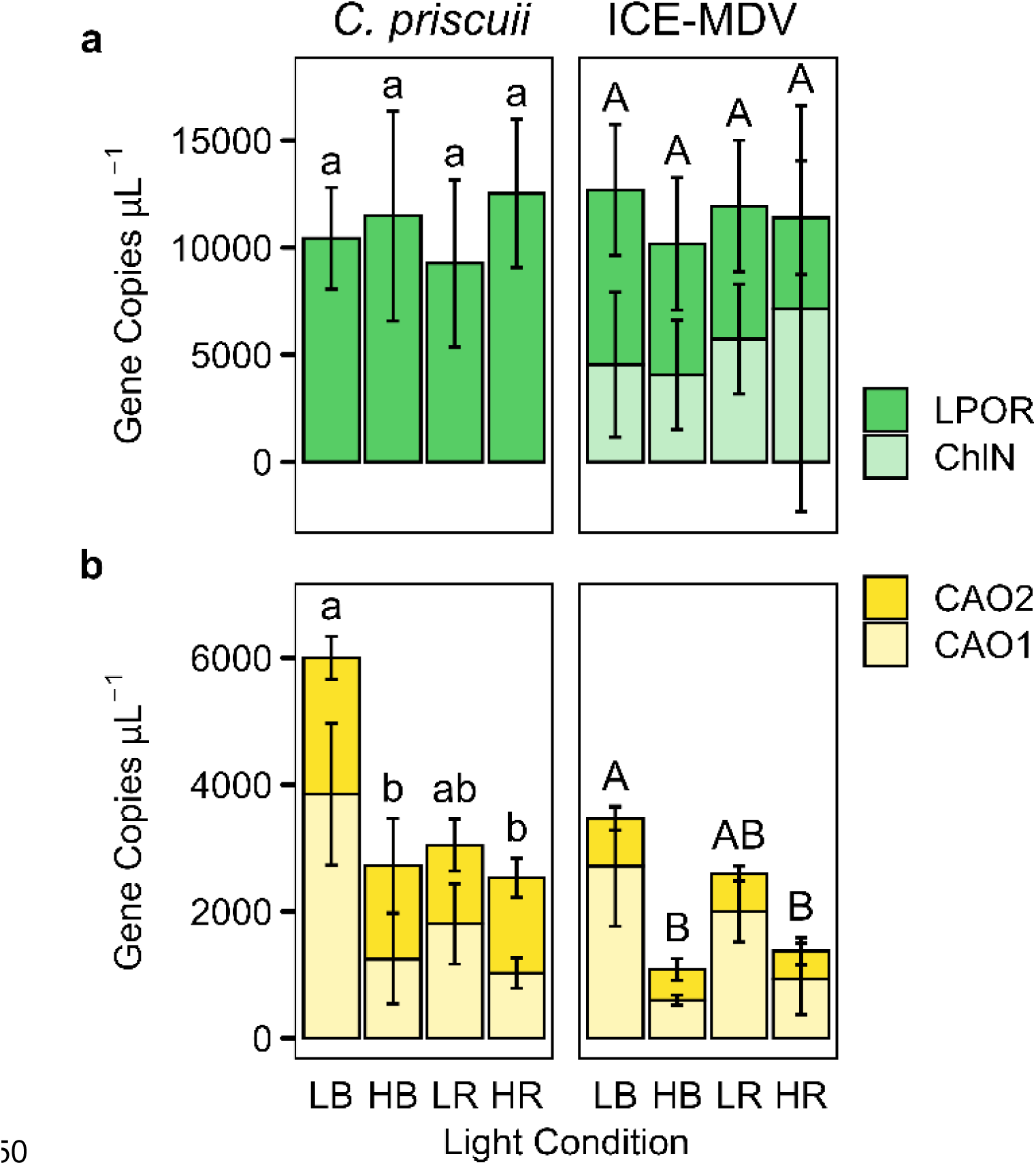
Transcript concentrations of LPOR and ChlN (a), and CAO1 and CAO2 (b) in *C. priscuii* and ICE-MDV grown in steady state light conditions measured using ddPCR. In both *C. priscuii* and ICE-MDV CAO1 expression was significantly affected by light quantity (two-way ANOVA, p=0.004 and p=0.002, respectively) while only *C. priscuii* was significantly affected by light quality (two-way ANOVA, p=0.03). CAO2 expression was not impacted by light quality or quantity in either species (two-way ANOVA, p>0.05).Different letters above the bars represent statistically significant differences between light conditions as determined using Tukey’s post hoc test (p < 0.05). Lowercase letters indicate significant differences in *C. priscuii,* capital letters indicate significant differences in ICE-MDV. LB, low blue; HB, high blue; LR, low red; HR, high red. Values are means ± SD (n = 3)

The overall expression levels of CAO are affected by light treatment, with significant differences between the two gene homologs (Fig. 5b). In both species, the expression of CAO1 is significantly affected by light quantity. In *C. priscuii* (but not ICE-MDV) CAO1 expression is also significantly affected by light quality. In contrast, in both species CAO2 is expressed constitutively and neither light quality nor quantity have a significant effect on transcript levels. Overall, total CAO expression is higher in *C. priscuii* than ICE-MDV, with the highest levels observed in algae acclimated to low blue light (∼6000 copies/µL vs ∼3400 copies/µL, respectively). Thus, it appears that the expression of CAO1 (but not CAO2) correlates with the ability of these species to modify Chl *a/b* ratios in response to light (Fig. 3b).

### Light-mediated changes in thylakoid polypeptide abundance

Differences in light quality and abundance are typically accompanied by long-term adjustments in PSI/PSII stoichiometry and PSII antenna size (Aizawa et al. 1992; Ballottari et al. 2007; Bonente et al. 2012; Melis et al. 1996). To investigate whether the abundance of major thylakoid proteins in the psychrophiles is affected by light, we analyzed the relative protein accumulation of PsaA, PsbA, and Lhcbm in cultures acclimated to different light conditions (Fig. 6). PsbA accumulation did not respond to light treatment in either species. In both species, acclimation to higher intensity light leads to decreases in PsaA and Lhcbm levels, indicating an increase in the size of the PSII antenna and decrease in PSI/PSII stoichiometry. This suggests that both psychrophiles retain the ability to regulate the size of their light harvesting apparatus when exposed to higher intensity light, but this ability is not linked to the spectral composition of the available light.

**Fig. 6.**
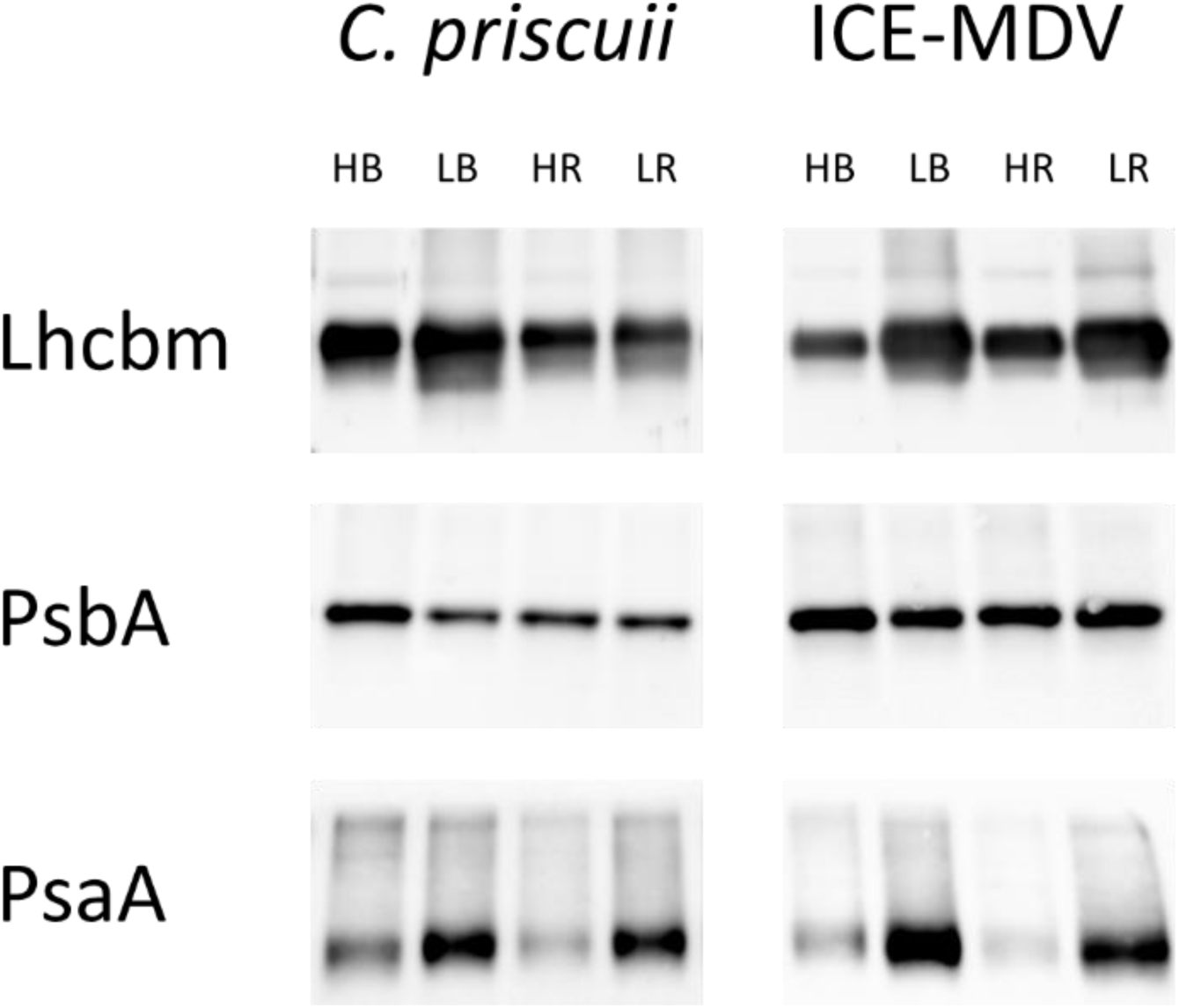
Representative immunoblots (n = 3) showing accumulation of Lhcbm, PsbA, and PsaA protein in *C. priscuii* and ICE-MDV in varied light treatments. Lanes were equally loaded with 10 µg of total protein. HB, high blue; LB, low blue; HR, high red; LR, low red

We also determined that the genomes of *C. priscuii* and ICE-MDV encode for a full complement of LHC genes, comparable in number to the model *C. reinhardtii* (Table S3). The genes encoding for LHCA proteins share a close relationship to their mesophilic counterparts, but the genes encoding for the major classes of LHCB proteins (LHCBMs) are more similar to their closest homologs within species rather than to their inter-species counterparts (Fig. S4).

### Light-driven differences in PSII photochemistry between *C. priscuii* and ICE-MDV

We observed major differences in total Chl amounts in *C. priscuii* and ICE-MDV grown under blue and red light (Fig. 3). To investigate whether differing Chl amounts affect the photosynthetic ability of these species, we measured Chl *a* fluorescence at the level of PSII. *C. priscuii* has very low maximum PSII photochemical efficiency (F_V_/F_M_) in higher intensity red light while ICE-MDV has moderately decreased levels of F_V_/F_M_ (0.15 and 0.34, respectively), when compared to values previously reported for green algae in optimal conditions (Bonente et al. 2012; Qin et al. 2021). We observed similar F_V_/F_M_ values in the cultures acclimated to blue or lower intensity red light (∼0.6-0.7; Fig. 7). A low F_V_/F_M_ can be interpreted as a damage to the photosynthetic machinery (Qin et al. 2021), suggesting more severe photoinhibition at the level of PSII in *C. priscuii* compared to ICE-MDV in higher red light (Fig. 7). *C. priscuii* acclimated to red light had very low Chl levels, regardless of the intensity (Fig. 3) but we observed low F_V_/F_M_ values only in higher intensity red light (Fig. 7). These results indicate that reduced capacity for PSII photochemistry observed in *C. priscuii* in high red light is not caused by low Chl levels.

**Fig. 7.**
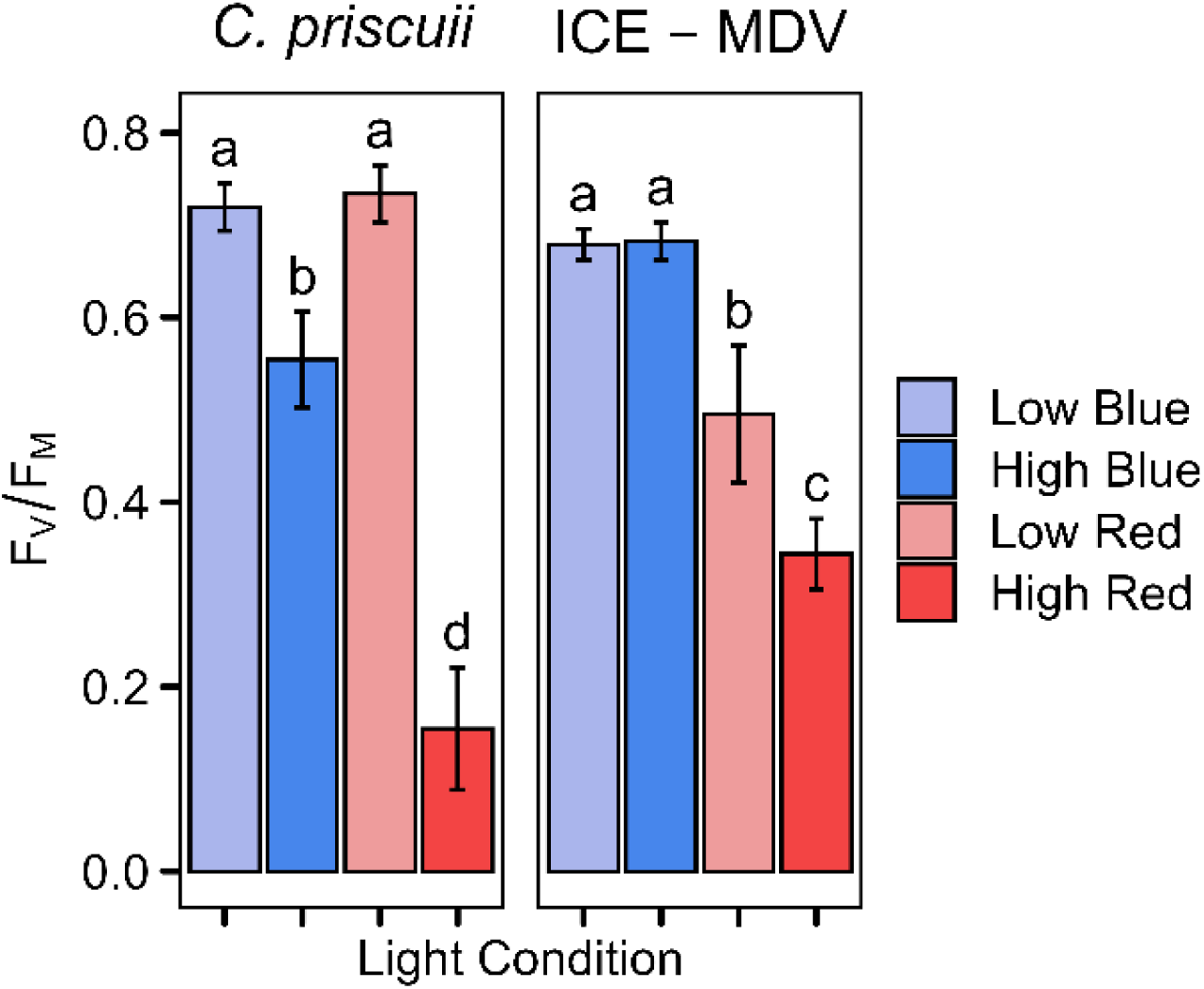
Maximum PSII photochemical efficiency (F_V_/F_M_) in *C. priscuii* and ICE-MDV cultures grown in different steady state light conditions. Statistically significant differences are represented as different letters as determined using Tukey’s post hoc test (p < 0.05). Values are means ± SD (n ≥ 3)

Light energy absorbed by PSII can be used for 1) photochemistry [Y(II)], 2) dissipated through regulated processes and antenna quenching [Y(NPQ)], or 3) through non-regulated processes [Y(NO)]; our results demonstrate a very different light partitioning mechanisms in the two psychrophiles. *C. priscuii* grown in higher intensity red light has a very low proportion of absorbed light energy used for photochemistry (Fig. 8a) or dissipated as NPQ (Fig. 8b; Fig. S5). Our data demonstrates that the largest proportion of absorbed light energy is dissipated in non-regulated ways as Y(NO), even at very low actinic light levels (Fig. 8c). This effect appears to be limited only to cultures acclimated to higher intensity red light, and cultures acclimated to blue or lower intensity red light maintain the capacity to use light energy through photochemistry or dissipate the excess in regulated ways as Y(NPQ). In contrast, ICE-MDV grown in higher intensity red light uses a larger proportion of absorbed light for photochemistry (Fig. 8d) and can effectively dissipate excess energy through NPQ (Fig. 8e; Fig. S5). ICE-MDV is also able to maintain a similar proportion of Y(NO) in all tested light conditions (Fig. 8f). We also examined NPQ as a measure independent of light partitioning as Y(NPQ) and observed the same trend (Fig. S5). Finally, we observed rapid NPQ relaxation kinetics in the dark in all cases, even when the NPQ values in the light were very high (Fig. S6), indicating that the majority of NPQ was reversible energy-dependent quenching and not photoinhibitory quenching (Ralph and Gademann 2005).

**Fig. 8.**
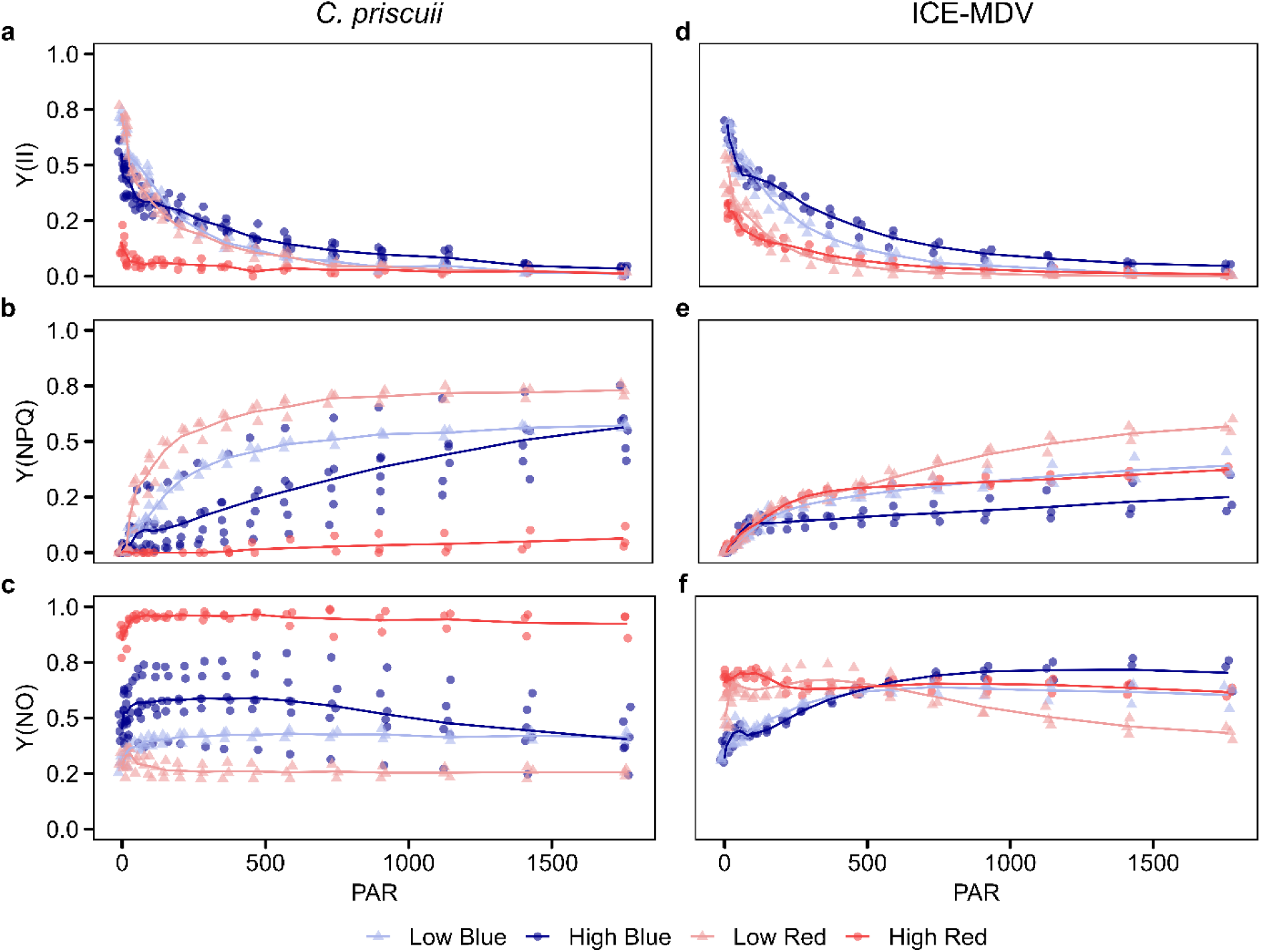
Effect of varied light conditions on energy partitioning in PSII in *C. priscuii* (a, b, c) and ICE-MDV (d, e, f). Total absorbed light distributed between: photochemical yield of PSII, Y(II); nonphotochemical quenching, Y(NPQ); nonregulated energy dissipation, Y(NO). Points represent individual biological replicates and lines represent mean (n ≥ 3)

We also measured the saturation point (Ek), utilization efficiency of light energy (α), and maximum rates of electron transport (rETR_max_). Both species have comparable saturation points in higher intensity red and blue light, suggesting similar light intensity tolerance (Jauffrais et al. 2022; Qin et al. 2021). Similarly, both species have reduced light utilization efficiencies and maximum rates of electron transport when grown in higher intensity red light, but this reduction is significantly more pronounced in *C. priscuii* compared to ICE-MDV (Table 1). This indicates a much-reduced ability to use higher intensity red light and a reduced ability to move electrons through the photosynthetic apparatus under these conditions, particularly for *C. priscuii* (Table 1). Taken together, our photosynthetic measurements suggest that *C. priscuii* has significantly reduced ability for efficient photochemistry when grown in red light. Excess energy is dissipated as NPQ when light intensity is low but predominantly as NO when the cells are acclimated to higher intensity red light. These effects are much less pronounced in ICE-MDV.

**Table 1.** Photosynthetic parameters derived from fitting curves of relative electron transport rate during rapid light curves in cultures of *C. priscuii* and ICE-MDV grown in different steady state light conditions. Light harvesting efficiency, α; light-saturated electron transport rate, ETR_max_; minimum saturating irradiance, Ek. Statistically significant differences are represented as different letters as determined using Tukey’s post hoc test (p < 0.05). LB, low blue; HB, high blue; LR, low red; HR, high red. Values are means ± SD (n ≥ 3)

| | $\alpha$ | | $E_k$ | | $rETR_{max}$ | |
| --- | --- | --- | --- | --- | --- | --- |
|  | *C. priscuii* | ICE-MDV | *C. priscuii* | ICE-MDV | *C. priscuii* | ICE-MDV |
| LB | $0.28 \pm 0.03^a$ | $0.28 \pm 0.03^{ab}$ | $88.0 \pm 16.8^b$ | $106.3 \pm 18.4^b$ | $24.5 \pm 3.5^b$ | $29.0 \pm 1.6^b$ |
| HB | $0.17 \pm 0.04^c$ | $0.26 \pm 0.03^{ab}$ | $254.7 \pm 129.8^a$ | $170.6 \pm 33.9^{ab}$ | $39.7 \pm 10.6^a$ | $43.9 \pm 6.7^a$ |
| LR | $0.23 \pm 0.06^b$ | $0.16 \pm 0.02^c$ | $99.9 \pm 29.8^b$ | $70.0 \pm 20.4^b$ | $21.4 \pm 3.7^{bc}$ | $11.8 \pm 4.3^d$ |
| HR | $0.04 \pm 0.01^e$ | $0.11 \pm 0.02^d$ | $271.1 \pm 137.9^a$ | $138.4 \pm 41.5^b$ | $9.0 \pm 2.9^d$ | $14.4 \pm 3.6^{cd}$ |

## Discussion

### Chlorophyll biosynthesis in Lake Bonney: the mystery deepens

While Chl is an essential and abundant pigment in all photosynthetic species but its biosynthesis requires tight regulation (Kobayashi and Masuda 2019; Tanaka and Tanaka 2007). Confirming previous computational analyses (Cvetkovska et al. 2018a) we experimentally show that DPOR is absent in *C. priscuii* (Fig. 4); and with it presumably the ability to synthesize Chl in the absence of light. As predicted (Smith et al. 2019), the genes encoding for the DPOR subunits are present in ICE-MDV (Fig. 4). The loss of DPOR in *C. priscuii* is not easily explained, since the presence of a light-independent enzyme in the light-limited environments of Lake Bonney may be seen as a benefit for maintaining adequate Chl levels. Indeed, it has been suggested that DPOR is particularly effective in deep or turbid aquatic environments where light limitations may pose a challenge for maintaining LPOR activity (Hunsperger et al. 2015).

While DPOR has been independently lost throughout eukaryotic evolution, most notably in all angiosperms, many algal species have retained the genes encoding for the subunits of this enzyme (Fong and Archibald 2008; Hunsperger et al. 2015; Wicke et al. 2011). The presence of both DPOR and LPOR in ICE-MDV conforms to this pattern, as this alga appears well equipped to modify its cellular Chl levels in the light conditions tested here (Fig. 3a). The endolithic coral symbiont *Ostreobium quekettii* (Bryopsidales) is currently the only confirmed eukaryote that depends solely on DPOR for Chl biosynthesis, as detailed screening of its genome failed to detect the LPOR gene (Iha et al. 2021; Marcelino et al. 2016). The authors surmise that reliance on DPOR may be advantageous due to the extreme shading this alga experiences within the coral skeleton but *C. priscuii* appears to have developed different strategies for shade adaptation.

In some species, LPOR duplication may compensate for the loss of DPOR (Hunsperger et al. 2015) but this does not appear to be the case for *C. priscuii*. Despite having a genome enriched in gene duplicates (Zhang et al. 2021), a second copy of LPOR was not detected by Cvetkovska et al. (2018a) or in this study. Instead, our results suggest that this alga compensates for the lack of DPOR by increasing LPOR expression levels (Fig. 5a). The regulation of LPOR expression has been extensively studied in angiosperms and involves a complex network of developmental, hormonal, and light signals (Gabruk and Mysliwa-Kurdziel 2015; Kobayashi and Masuda 2019; Yong et al. 2024) but the situation is less clear in algae. While phytohormones have been detected in algae, their physiological roles remain unknown (Lu and Xu 2015). Phytochrome and cryptochrome photoreceptors are important regulators of LPOR expression in angiosperms, but *C. priscuii* lacks phytochromes (like all Chlorophytes; Li et al. 2015) and encodes for a significantly reduced complement of cryptochrome genes compared to its relatives (Poirier et al. 2023). The coordination of LPOR and DPOR expression in species that have both (such as ICE-MDV) is also poorly understood, although complementation of LPOR downregulation by DPOR has been reported in the cyanobacterium *Fremyella diplosiphon* (Shui et al. 2009).

Consequences of the dependence on the light-active LPOR may be seen in the attenuated ability of *C. priscuii* to synthesize Chl in response to light availability. This is particularly obvious in cultures grown in red light that accumulate very low Chl levels regardless of the intensity (Fig. 3a). Work with land plants has shown that the LPOR substrate Pchlide has absorbance maxima in both red and blue regions of the light spectrum (Koski et al. 1948), but that the conversion of Pchlide to Chlide by LPOR was up to seven times more efficient when Pchlide absorbed red light (647 nm) rather than blue light (407 nm) (Hanf et al. 2012). The predicted high activity of land plant LPOR in red light contrasts with the low Chl levels observed in red light in *C. priscuii*. This presents another puzzle regarding the role of LPOR in chlorophyll biosynthesis in *C. priscuii*, an alga that is exposed solely to blue light in its habitat. Although it must be noted that the validity of the results by Hanf et al. (2012) have been questioned (Björn 2013), there could be several other explanations to the low Chl levels observed in red light-grown *C. priscuii* cultures. First, the Chl biosynthesis pathway could be light constrained at an upstream step. For instance, the synthesis of 5-aminolevulinic acid (ALA), the universal precursor of tetrapyrrolesis also light regulated (Ilag et al. 1994; reviewed in Wu et al. 2019). Second, deep water algae such as *C. priscuii* may have a “blue-adapted” LPOR which is more efficient at shorter wavelengths. There are several reports of proteins from *C. priscuii* that are specifically adapted to function better in the extreme environment of Lake Bonney (Cvetkovska et al. 2018b; Szyszka-Mroz et al. 2019), demonstrating the possibility for evolving beneficial protein traits under environmental pressures. Detailed biochemical and transcriptional characterization of LPOR from psychrophilic algae would shed more light on the mystery of Chl biosynthesis in the depths of Lake Bonney and other blue light dominated environments.

### CAO gene duplication may confer a fine control of Chl *b* accumulation in polar algae

Both *C. priscuii* and ICE-MDV encode for two CAO genes, representing the first report of CAO duplication in algae. This duplication was reported previously based on genomic analysis in *C. priscuii* (Cvetkovska et al. 2018a), but here we provide experimental evidence that both Lake Bonney psychrophiles encode for two highly similar CAO homologs (Fig. S3). CAO duplication has been reported only twice. Rice (*Oryza sativa*) encodes for two CAO genes, although only CAO1 is functional and necessary for maintaining Chl *b* amounts (Jung et al. 2021; Lee et al. 2005). The bryophyte *Physcomitrium patens* also encodes for two CAO genes, both of which contribute to Chl *b* biosynthesis (Zhang et al. 2023). CAO is encoded by a single gene in all other examined plants and algae (Kunugi et al. 2016; Schumacher et al. 2022).

We believe that the CAO duplications arose independently in *C. priscuii* and ICE-MDV, possibly driven by the environmental pressures of their extreme environment. While both species were isolated from Lake Bonney, they are not the closest relatives within the Chlamydomonadales; *C. priscuii* belongs to the Moewusinia clade (Possmayer et al. 2016) while ICE-MDV (and related ICE strains) in the Monadinia clade (Cook et al. 2019; Zhang et al. 2020). Furthermore, the two CAO homologs share a closer identity within species (Fig. S3), suggesting an independent duplication event.

It has been hypothesized that CAO gene expression is the primary method for regulating chl *a/b* ratios and antenna size (Tanaka and Tanaka 2019). In both species, CAO2 is constitutively expressed, while CAO1 is upregulated in cells acclimated to lower light intensity (although the change is significant only in blue light). Higher expression of CAO under lower intensity light (Fig. 5b) correlates with increased Chl *b* pigment (Fig. 3b) and LHCBM protein (Fig. 6) amounts in both *C. priscuii* and ICE-MDV. Accumulation of Chl *b* affects the amount of LHCII present and in many plant and algal species growth in low light leads to higher Chl *b* amounts and larger antenna size (Harper et al. 2004; Kunugi et al. 2016; Masuda et al. 2003; Stolárik et al. 2017; Tanaka and Tanaka 2005) including in C*. priscuii* (Szyszka et al. 2007). It is tempting to speculate that CAO duplication in these polar species leads to increased CAO protein amounts through gene dosage, as demonstrated for photosynthetic ferredoxin (Cvetkovska et al. 2018b). This could support a finer control for increasing Chl *b* amounts and large PSII antennae in the light-limited environment of Lake Bonney. This would likely offer competitive advantage and be a very beneficial trait for maximizing light absorption under the polar ice.

We also observed similar duplication patterns in the genes encoding for LHCBM proteins. LHCBM gene duplications were reported previously in the genome of *C. priscuii* (Zhang et al. 2021), and here we show that both ICE-MDV and *C. priscuii* have more genes that encode for Type IV (LHCBM1) and Type III (LHCBM2/7) proteins but lack Type II (LHCBM5) and have a reduced complement of Type I (LHCBM3/4/6/8/9) genes (Fig. S4). LHCBM genes are also more similar within species rather than to their homologs from other species, suggesting independent gene evolution. Interestingly, this appears to be the case only for LHCBM genes, and both species encode for a full complement of LHCA genes that share a close identity between species including *C. reinhardtii* (Fig. S4). Inherently low levels of PSI and the failure to detect LHCA proteins using immunoblotting in *C. priscuii*, combined with its inability to preform state transitions (Morgan et al. 1998; Morgan-Kiss et al. 2005; Szyszka et al. 2007) has led to questioning the importance of PSI in this alga. The more recent characterization of a unique PSI-supercomplex, the discovery of energy spillover between PSI and PSII, and the detection of PSI peptides via proteomics (Kalra et al. 2020; Kalra et al. 2023; Szyzszka-Mroz et al. 2015; Szyszka-Mroz et al. 2019) revealed that PSI plays a different role in *C. priscuii* than its model relatives. Importantly, these studies also failed to detect a full complement of LHCBM peptides in *C. priscuii* (Kalra et al. 2020; Kalra et al. 2023), explained here by the lack of the genes encoding them. Previous studies along with the genetic data we present here are revealing a unique photosynthetic apparatus in shade-adapted psychrophiles.

### Red light negatively affects photosynthetic performance in *C. priscuii*

Up to this point, our results suggest that the two psychrophiles acclimated to different light conditions respond more strongly to light intensity rather than light quality, possibly related to decreased ability to sense and respond to spectral wavelengths (Poirier et al. 2023). These results contrast with those reported by Morgan-Kiss et al. (2005), where a sudden shift from white to red light caused a complete growth inhibition and led to re-arrangement in the stoichiometry of the two photosystems in *C. priscuii*. These differences in responses between a sudden shift and long-term growth under different spectral wavelengths prompted us to further examine the impact of light quality on photochemistry.

Higher intensity red light has a significant and negative effect on photochemistry in *C. priscuii* as demonstrated by very low F_V_/F_M_ levels (Fig. 7), an effect not observed with lower intensity red light nor blue light. Higher intensity red light also dramatically affects light partitioning in this alga, and we observe most of the light energy being dissipated through non-regulated processes as Y(NO) rather than non-photochemical quenching as Y(NPQ) (Fig. 8). The light saturation coefficient (Ek) can be used as a proxy for the ability to tolerate higher light (Jauffrais et al. 2022; Qin et al. 2021; Ralph and Gademann 2005). Comparable Ek values in high red light to those in blue light suggests that *C. priscuii* cells are absorbing similar light levels; however, in high red light they are not able to utilize this light for photochemistry or dissipate it through NPQ. Significantly, this effect is observed solely in algae grown at higher intensity red light, and higher intensity blue light does not appear to have the same photoinhibitory effect. This suggests that photoacclimation methods such as alteration of antenna size, PSI/PSII stoichiometry, Chl levels, and induction of NPQ are sufficient to avoid photoinhibition in blue light (even at higher intensity) but not in red light.

Previous work has shown that *C. priscuii* has a higher capacity for NPQ, compared to related algae. For instance, when grown at optimal temperatures Y(NPQ) was almost twice as high in *C. priscuii* compared to the related mesophile *Chlamydomonas raudensis* SAG 49.72 (Szyszka et al., 2007). Growth at higher intensity white light led to doubling of both Y(NPQ) and Y(NO) when compared to growth at lower light (Szyszka et al. 2007) and an increase of NPQ was reported when cultures were rapidly shifted from white to red light (Morgan-Kiss et al. 2005). High Y(NO) and low Y(NPQ), reminiscent to those seen in our high red light grown cultures, were reported in *C. priscuii* multi-cell palmelloids (Szyszka-Mroz et al. 2022). The authors suggested that the formation of palmelloids is a photoprotective mechanism in *C. priscuii,* but it is unlikely that a similar process is occurring in our red light-growth cultures. Palmelloids are larger cellular assemblages (Szyszka-Mroz et al. 2022), and we did not observe an increase in average cell size in red-grown cells when compared to those grown in blue light (Fig. S1). Thus, it appears that very low NPQ rates are unique to high red light acclimated *C. priscuii*.

While red light also decreases photochemistry in ICE-MDV, the effect is much less pronounced. ICE-MDV grown at higher intensity red light has moderately decreased F_V_/F_M_ values (∼0.34) and slightly increased but comparable levels of Y(NPQ) and Y(NO) as blue light-grown cultures. Unlike *C. priscuii*, ICE-MDV has retained the ability to modify its photosynthetic apparatus under Fe-deficiency (Cook et al. 2019), can balance PSI and PSII light excitation through state transitions (Kalra et al. 2023) and responds phototactically to light signals (Poirier et al. 2023) suggesting a photosynthetic apparatus similar to the mesophile *C. reinhardtii*. ICE-MDV is naturally adapted to a wider range of light conditions in Lake Bonney (Li and Morgan-Kiss 2019; Sherwell et al. 2022) compared to *C. priscuii,* which is reflected in its ability to maintain photosynthetic efficiencies under both red and blue light.

One of the more puzzling findings of our study is the observation that *C. priscuii* can maintain relatively high growth rates and low cellular death in higher intensity red light (Fig. 2), despite very low Chl levels (Fig. 3) and severely inhibited photosynthetic capacity (Fig. 7, Fig. 8, Table 1). The very low F_V_/F_M_ values in this alga acclimated to red light (∼0.15) are reminiscent of polar microalgae that undergo very low growth rates or dormancy during extended periods of darkness (∼0.1; Reeves et al. 2011; McMinn and Hegseth 2004). Many algal species, including *C. reinhardtii* and various polar alga, are mixotrophs and can use external carbon sources to support growth in the absence of light (Heifetz et al. 2000; Stoecker et al. 2018). All our experiments were performed under photoautotrophic conditions, so it is very unlikely that our cultures were using external organic compounds to fuel growth. One possibility is that most of the red light is preferentially absorbed by PSI (Melis et al. 1996), a parameter not measured by the methods used in our work. The importance of PSI and CEF within a unique supercomplex in *C. priscuii* has received significant attention recently (Kalra et al. 2020; Kalra et al. 2023; Szyszka-Mroz et al. 2015), but all these studies have utilized a broad range white light. Future work examining the involvement of CEF and PSI under narrow-wavelength growth will shed further light on the unique photochemistry in deep-water psychrophiles.

## Conclusion

Overall, we suggest that the unique physiology of *C. priscuii* is best suited for the extreme but very stable environment of Lake Bonney, Antarctica. Many psychrophilic traits that may confer adaptive benefits for life at the extreme, could be detrimental when the environmental conditions change (Cvetkovska et al. 2022a). Thus, the loss of short-term acclimation mechanisms may be what limits *C. priscuii* to the deep photic zone of Lake Bonney and what may ultimately harm its viability in a changing world. In contrast, ICE-MDV is much better equipped to manage changes in the spectral composition of light. Rising temperatures are predicted to alter light availability within Lake Bonney as the ice cover thins, increasing the light intensity and variation in light quality (Clark et al. 2013; Howard-Williams et al. 1998). Reduced photochemistry and growth due to changes in spectral composition would likely be a disadvantage in a competitive natural environment. We used narrow-range wavelengths to better understand the effects of light quality, but observing the responses of these vulnerable species to white light enriched in red and blue wavelengths would be the next step to better understand the responses of these psychrophiles to a changing world.

## Statements and Declarations

### Data availability statement

The genomic data that support the findings of this study are available in Phytozome (https://phytozome-next.jgi.doe.gov/) and NCBI GenBank (https://www.ncbi.nlm.nih.gov/genbank/). All accession numbers for the sequences are available within the Supplementary Data (Supplementary Table S2 and S3). All other data that support the findings of this study are available from the corresponding author upon reasonable request.

## Supplemental data

**Fig. S1** Average size of *C. priscuii* and ICE-MDV cells grown in different steady state light conditions

**Fig. S2** Alignment of LPOR sequences from representative chlorophyte species

**Fig. S3** Multiple alignment of CAO peptides from *C. priscuii*, ICE-MDV and representative green algae.

**Fig. S4** Phylogenetic analysis of LHCA (a) and LHCBM (b) genes from *C. priscuii*, ICE-MDV and *C. reinhardtii*

**Fig. S5** Nonphotochemical quenching (NPQ) at increasing light intensity measured during rapid light curve in *C. priscuii* and ICE-MDV cultures grown in different steady state light conditions

**Fig. S6** NPQ rates during rapid light curve and dark recovery effects over time in *C. priscuii* and ICE-MDV cultures grown in different steady state light conditions

**Table S1** Sequences of species-specific primers used to measure CAO, LPOR, and CHLN gene expression and conserved primers used to confirm presence or absence of DPOR subunits in this study

**Table S2** Genes involved in chlorophyll biosynthesis encoded in the genomes of *Chlamydomonas priscuii* and *Chlamydomonas* sp. ICE

**Table S3** Light Harvesting Complex (LHC) genes encoded in the genomes of *Chlamydomonas priscuii* and *Chlamydomonas* sp. ICE

## Author contributions

M.C. and M.P. conceptualized the work and designed the experiments. M.P. preformed the majority of experiments and wrote the initial draft of the manuscript. K.F. contributed to the bioinformatics analysis of LHC genes and M.C. contributed to bioinformatic analysis of chlorophyll biosynthesis genes. M.C. supervised the work. All authors contributed towards manuscript preparation and editing.

## Acknowledgements

The authors wish to thank Dr. James Raymond (University of Nevada, NV, USA) and Dr. Rachael Morgan-Kiss (Miami University, OH, USA) for providing access and assistance with the ICE-MDV genomic data. We would also like to thank the Faculty of Science and the Core Facilities Program at uOttawa for instrument access and support.

## References

1. Aizawa, K., Shimizu, T., Hiyama, T., Satoh, K., and Nakamura, Y. (1992). Changes in composition of membrane proteins accompanying the regulation of PS I/PS II stoichiometry observed with *Synechocystis* PCC 6803. Photosynthesis Research, 32(2), 131–138. 10.1007/BF00035947

2. Allorent, G., and Petroutsos, D. (2017). Photoreceptor-dependent regulation of photoprotection. Current Opinion in Plant Biology, 37, 102–108. 10.1016/j.pbi.2017.03.016

3. Armaleo, D., Müller, O., Lutzoni, F., Andrésson, Ó. S., Blanc, G., Bode, H. B., … Xavier, B. B. (2019). The lichen symbiosis re-viewed through the genomes of Cladonia grayi and its algal partner Asterochloris glomerata. BMC Genomics, 20(1), 1–33. 10.1186/s12864-019-5629-x

4. Ballottari, M., Dall’Osto, L., Morosinotto, T., and Bassi, R. (2007). Contrasting behavior of higher plant photosystem I and II antenna systems during acclimation. Journal of Biological Chemistry, 282(12), 8947–8958. 10.1074/jbc.M606417200

5. Bielewicz, S., Bell, E., Kong, W., Friedberg, I., Priscu, J. C., and Morgan-Kiss, R. M. (2011). Protist diversity in a permanently ice-covered Antarctic Lake during the polar night transition. ISME Journal, 5(9), 1559–1564. 10.1038/ismej.2011.23

6. Biswal, A. K., Pattanayak, G. K., Pandey, S. S., Leelavathi, S., Reddy, V. S., Govindjee, and Tripathy, B. C. (2012). Light intensity-dependent modulation of chlorophyll b biosynthesis and photosynthesis by overexpression of chlorophyllide a oxygenase in tobacco. Plant Physiology, 159(1), 433–449. 10.1104/pp.112.195859

7. Biswal, A. K., Pattanayak, G. K., Ruhil, K., Kandoi, D., Mohanty, S. S., Leelavati, S., … Tripathy, B. C. (2024). Reduced expression of chlorophyllide a oxygenase (CAO) decreases the metabolic flux for chlorophyll synthesis and downregulates photosynthesis in tobacco plants. Physiology and Molecular Biology of Plants, 30(1), 1–16. 10.1007/s12298-023-01395-5

8. Björn, L. O. (2013). Comment on “Catalytic efficiency of a photoenzyme - An adaptation to natural light conditions” by J. Popp et al. ChemPhysChem, 14(11), 2595–2597. 10.1002/cphc.201300082

9. Blanc, G., Agarkova, I., Grimwood, J., Kuo, A., Brueggeman, A., Dunigan, D. D., … Van Etten, J. L. (2012). The genome of the polar eukaryotic microalga Coccomyxa subellispsoidea reveals traits of cold adaptation. Genome Biology, 13(1), 1–12. 10.2216/i0031-8884-4-1-43.1

10. Blanc, G., Duncan, G., Agarkova, I., Borodovsky, M., Gurnon, J., Kuo, A., … van Etten, J. L. (2010). The Chlorella variabilis NC64A genome reveals adaptation to photosymbiosis, coevolution with viruses, and cryptic sex. Plant Cell, 22(9), 2943–2955. 10.1105/tpc.110.076406

11. Bogen, C., Al-Dilaimi, A., Albersmeier, A., Wichmann, J., Grundmann, M., Rupp, O., … Kruse, O. (2013). Reconstruction of the lipid metabolism for the microalga Monoraphidium neglectum from its genome sequence reveals characteristics suitable for biofuel production. BMC Genomics, 14(1). 10.1186/1471-2164-14-926

12. Bonente, G., Pippa, S., Castellano, S., Bassi, R., and Ballottari, M. (2012). Acclimation of Chlamydomonas reinhardtii to different growth irradiances. Journal of Biological Chemistry, 287(8), 5833–5847. 10.1074/jbc.M111.304279

13. Bujaldon, S., Kodama, N., Rappaport, F., Subramanyam, R., de Vitry, C., Takahashi, Y., and Wollman, F. A. (2017). Functional Accumulation of Antenna Proteins in Chlorophyll b-Less Mutants of Chlamydomonas reinhardtii. Molecular Plant, 10(1), 115–130. 10.1016/j.molp.2016.10.001

14. Clark, G. F., Stark, J. S., Johnston, E. L., Runcie, J. W., Goldsworthy, P. M., Raymond, B., and Riddle, M. J. (2013). Light-driven tipping points in polar ecosystems. Global Change Biology, 19(12), 3749–3761. 10.1111/gcb.12337

15. Cook, G., Teufel, A., Kalra, I., Li, W., Wang, X., Priscu, J., and Morgan-Kiss, R. (2019). The Antarctic psychrophiles Chlamydomonas spp. UWO241 and ICE-MDV exhibit differential restructuring of photosystem I in response to iron. Photosynthesis Research, 141(2), 209–228. 10.1007/s11120-019-00621-0

16. Craig, R. J., Gallaher, S. D., Shu, S., Salomé, P. A., Jenkins, J. W., Blaby-Haas, C. E., … Merchant, S. S. (2023). The Chlamydomonas Genome Project, version 6: Reference assemblies for mating-type plus and minus strains reveal extensive structural mutation in the laboratory. Plant Cell, 35(2), 644–672. 10.1093/plcell/koac347

17. Cvetkovska, M., Orgnero, S., Hüner, N. P. A., and Smith, D. R. (2018a). The enigmatic loss of light-independent chlorophyll biosynthesis from an Antarctic green alga in a light-limited environment. New Phytologist, 222(2). 10.1111/nph.15623

18. Cvetkovska, M., Szyszka-Mroz, B., Possmayer, M., Pittock, P., Lajoie, G., Smith, D. R., and Hüner, N. P. A. (2018b). Characterization of photosynthetic ferredoxin from the Antarctic alga Chlamydomonas sp. UWO241 reveals novel features of cold adaptation. New Phytologist, 219(2), 588–604. 10.1111/nph.15194

19. Cvetkovska, M., Vakulenko, G., Smith, D. R., Zhang, X., and Hüner, N. P. A. (2022a). Temperature stress in psychrophilic green microalgae: Minireview. Physiologia Plantarum, 174(6). 10.1111/ppl.13811

20. Cvetkovska, M., Zhang, X., Vakulenko, G., Benzaquen, S., Szyszka-Mroz, B., Malczewski, N., … Hüner, N. P. A. (2022b). A constitutive stress response is a result of low temperature growth in the Antarctic green alga Chlamydomonas sp. UWO241. Plant Cell and Environment, 45(1), 156–177. 10.1111/pce.14203

21. da Roza, P. A., Muller, H., Sullivan, G. J., Walker, R. S. K., Goold, H. D., Willows, R. D., … Paulsen, I. T. (2024). Chromosome-scale assembly of the streamlined picoeukaryote Picochlorum sp. SENEW3 genome reveals Rabl-like chromatin structure and potential for C4 photosynthesis. Microbial Genomics, 10(4), 1–18. 10.1099/mgen.0.001223

22. De Maayer, P., Anderson, D., Cary, C., and Cowan, D. A. (2014). Some like it cold: Understanding the survival strategies of psychrophiles. EMBO Reports, 15(5), 508–517. 10.1002/embr.201338170

23. Dolhi, J. M., Maxwell, D. P., and Morgan-Kiss, R. M. (2013). Review: The Antarctic Chlamydomonas raudensis: an emerging model for cold adaptation of photosynthesis. Extremophiles, 17(5), 711– 722. 10.1007/s00792-013-0571-3

24. Erickson, E., Wakao, S., and Niyogi, K. K. (2015). Light stress and photoprotection in Chlamydomonas reinhardtii. Plant Journal, 82(3), 449–465. 10.1111/tpj.12825

25. Fong, A., and Archibald, J. M. (2008). Evolutionary dynamics of light-independent protochlorophyllide oxidoreductase genes in the secondary plastids of cryptophyte algae. Eukaryotic Cell, 7(3), 550–553. 10.1128/EC.00396-07

26. Gabruk, M., and Mysliwa-Kurdziel, B. (2015). Light-Dependent Protochlorophyllide Oxidoreductase: Phylogeny, Regulation, and Catalytic Properties. Biochemistry, 54(34), 5255–5262. 10.1021/acs.biochem.5b00704

27. Goodstein, D. M., Shu, S., Howson, R., Neupane, R., Hayes, R. D., Fazo, J., … Rokhsar, D. S. (2012). Phytozome: A comparative platform for green plant genomics. Nucleic Acids Research, 40(D1), 1178–1186. 10.1093/nar/gkr944

28. Gooseff, M. N., Barrett, J. E., Adams, B. J., Doran, P. T., Fountain, A. G., Lyons, W. B., … Wall, D. H. (2017). Decadal ecosystem response to an anomalous melt season in a polar desert in Antarctica. Nature Ecology and Evolution, 1(9), 1334–1338. 10.1038/s41559-017-0253-0

29. Griffiths, W. T., McHugh, T., and Blankenship, R. E. (1996). The light intensity dependence of protochlorophyllide photoconversion and its significance to the catalytic mechanism of protochlorophyllide reductase. FEBS Letters, 398(2–3), 235–238. 10.1016/S0014-5793(96)01249-5

30. Grigoriev, I. V., Hayes, R. D., Calhoun, S., Kamel, B., Wang, A., Ahrendt, S., … Kuo, A. (2021). PhycoCosm, a comparative algal genomics resource. Nucleic Acids Research, 49(D1), D1004–D1011. 10.1093/nar/gkaa898

31. Hanf, R., Fey, S., Schmitt, M., Hermann, G., Dietzek, B., and Popp, J. (2012). Catalytic efficiency of a photoenzyme-An adaptation to natural light conditions. ChemPhysChem, 13(8), 2013–2015. 10.1002/cphc.201200194

32. Hanschen, E. R., Marriage, T. N., Ferris, P. J., Hamaji, T., Toyoda, A., Fujiyama, A., … Olson, B. J. S. C. (2016). The Gonium pectorale genome demonstrates co-option of cell cycle regulation during the evolution of multicellularity. Nature Communications, 7, 1–10. 10.1038/ncomms11370

33. Harper, A. L., Von Gesjen, S. E., Linford, A. S., Peterson, M. P., Faircloth, R. S., Thissen, M. M., and Brusslan, J. A. (2004). Chlorophyllide a oxygenase mRNA and protein levels correlate with the chlorophyll a/b ratio in Arabidopsis thaliana. Photosynthesis Research, 79(2), 149–159. 10.1023/B:PRES.0000015375.40167.76

34. Heifetz, P. B., Förster, B., Osmond, C. B., Giles, L. J., and Boynton, J. E. (2000). Effects of acetate on facultative autotrophy in Chlamydomonas reinhardtii assessed by photosynthetic measurements and stable isotope analyses. Plant Physiology, 122(4), 1439–1445. 10.1104/pp.122.4.1439

35. Hirooka, S., Hirose, Y., Kanesaki, Y., Higuchi, S., Fujiwara, T., Onuma, R., … Miyagishima, S. Y. (2017). Acidophilic green algal genome provides insights into adaptation to an acidic environment. Proceedings of the National Academy of Sciences of the United States of America, 114(39), E8304– E8313. 10.1073/pnas.1707072114

36. Howard-williams, C., Schwarz, A., Hawes, I., and Priscu, J. C. (1998). Optical properties of the McMurdo Dry Valley lakes, Antarctica. In Ecosystem Dynamics in a Polar Desert: the McMurdo Dry Valleys, Antarctica (pp. 189–203).

37. Hui, C., Schmollinger, S., and Glaesener, A. G. (2023). Chapter 11: Growth techniques. In The Chlamydomonas Sourcebook: Volume 1: Introduction to Chlamydomonas and Its Laboratory Use (Vol. 1). 10.1016/B978-0-12-822457-1.00005-4

38. Hüner, N. P. A., Smith, D. R., Cvetkovska, M., Zhang, X., Ivanov, A. G., Szyszka-Mroz, B., … Morgan-Kiss, R. (2022). Photosynthetic adaptation to polar life: Energy balance, photoprotection and genetic redundancy. Journal of Plant Physiology, 268(September 2021), 153557. 10.1016/j.jplph.2021.153557

39. Hunsperger, H. M., Randhawa, T., and Cattolico, R. A. (2015). Extensive horizontal gene transfer, duplication, and loss of chlorophyll synthesis genes in the algae. BMC Evolutionary Biology, 15(1), 1–19. 10.1186/s12862-015-0286-4

40. Iha, C., Dougan, K. E., Varela, J. A., Avila, V., Jackson, C. J., Bogaert, K. A., … Verbruggen, H. (2021). Genomic adaptations to an endolithic lifestyle in the coral-associated alga Ostreobium. Current Biology, 31(7), 1393–1402.e5. 10.1016/j.cub.2021.01.018

41. Ilag, L. L., Kumar, A. M., and Söll, D. (1994). Light regulation of chlorophyll biosynthesis at the level of 5-aminolevulinate formation in arabidopsis. Plant Cell, 6(2), 265–275. 10.2307/3869644

42. Jauffrais, T., Brisset, M., Lagourgue, L., Payri, C. E., Gobin, S., Le Gendre, R., and Van Wynsberge, S. (2022). Seasonal changes in the photophysiology of Ulva batuffolosa in a coastal barrier reef. Aquatic Botany, 179(February), 0–1. 10.1016/j.aquabot.2022.103515

43. Jeffrey, S. W., and Humphrey, G. F. (1975). New spectrophotometric equations for determining chlorophylls a, b, c1 and c2 in higher plants, algae and natural phytoplankton. Biochemie Und Physiologie Der Pflanzen, Vol. 167, pp. 191–194. 10.1016/s0015-3796(17)30778-3

44. Jung, Y. J., Lee, H. J., Yu, J., Bae, S., Cho, Y. G., and Kang, K. K. (2021). Transcriptomic and physiological analysis of OsCAO1 knockout lines using the CRISPR/Cas9 system in rice. Plant Cell Reports, 40(6), 1013–1024. 10.1007/s00299-020-02607-y

45. Kalra, I., Wang, X., Cvetkovska, M., Jeong, J., McHargue, W., Zhang, R., … Morgan-Kissa, R. (2020). Chlamydomonas sp. UWO 241 exhibits high cyclic electron flow and rewired metabolism under high salinity. Plant Physiology, 183(2), 588–601. 10.1104/pp.19.01280

46. Kalra, I., Wang, X., Zhang, R., and Morgan-Kiss, R. (2023). High salt-induced PSI-supercomplex is associated with high CEF and attenuation of state transitions. Photosynthesis Research, 157(2–3), 65–84. 10.1007/s11120-023-01032-y

47. Kim, E. H., Li, X. P., Razeghifard, R., Anderson, J. M., Niyogi, K. K., Pogson, B. J., and Chow, W. S. (2009). The multiple roles of light-harvesting chlorophyll a/b-protein complexes define structure and optimize function of Arabidopsis chloroplasts: A study using two chlorophyll b-less mutants. Biochimica et Biophysica Acta - Bioenergetics, 1787(8), 973–984. 10.1016/j.bbabio.2009.04.009

48. Kim, J. I., Moore, C. E., Archibald, J. M., Bhattacharya, D., Yi, G., Yoon, H. S., and Shin, W. (2017). Evolutionary dynamics of cryptophyte plastid genomes. Genome Biology and Evolution, 9(7), 1859– 1872. 10.1093/gbe/evx123

49. Kitajima, M., and Butler, W. L. (1975). Quenching of chlorophyll fluorescence and primary photochemistry in chloroplasts by dibromothymoquinone. BBA - Bioenergetics, 376(1), 105–115. 10.1016/0005-2728(75)90209-1

50. Kobayashi, K., and Masuda, T. (2019). Transcriptional control for the chlorophyll metabolism. In Advances in Botanical Research (1st ed., Vol. 91). 10.1016/bs.abr.2019.03.001

51. Koski, V. M., and Smith, J. H. C. (1948). The Isolation and Spectral Absorption Properties of Protochlorophyll from Etiolated Barley Seedlings. Journal of the American Chemical Society, 70(11), 3558–3562. 10.1021/ja01191a006

52. Kramer, D. M., Johnson, G., Kiirats, O., and Edwards, G. E. (2004). New fluorescence parameters for the determination of QA redox state and excitation energy fluxes. Photosynthesis Research, 79, 209– 218.

53. Król, M., Spangfort, M. D., Huner, N. P. A., Öquist, G., Gustafsson, P., and Jansson, S. (1995). Chlorophyll a/b-binding proteins, pigment conversions, and early light-induced proteins in a chlorophyll b-less Barley mutant. Plant Physiology, 107(3), 873–883. 10.1104/pp.107.3.873

54. Kumar, S., Stecher, G., Li, M., Knyaz, C., and Tamura, K. (2018). MEGA X: Molecular evolutionary genetics analysis across computing platforms. Molecular Biology and Evolution, 35(6), 1547–1549. 10.1093/molbev/msy096

55. Kunugi, M., Satoh, S., Ihara, K., Shibata, K., Yamagishi, Y., Kogame, K., … Tanaka, A. (2016). Evolution of Green Plants Accompanied Changes in Light-Harvesting Systems. Plant and Cell Physiology, 57(6), 1231–1243. 10.1093/pcp/pcw071

56. Lee, S., Kim, J. H., Eun, S. Y., Lee, C. H., Hirochika, H., and An, G. (2005). Differential regulation of chlorophyll a oxygenase genes in rice. Plant Molecular Biology, 57(6), 805–818. 10.1007/s11103-005-2066-9

57. Letunic, I., and Bork, P. (2024). Interactive Tree of Life (iTOL) v6: recent updates to the phylogenetic tree display and annotation tool. Nucleic Acids Research, 1–5. 10.1093/nar/gkae268

58. Li, F. W., Melkonian, M., Rothfels, C. J., Villarreal, J. C., Stevenson, D. W., Graham, S. W., … Mathews, S. (2015). Phytochrome diversity in green plants and the origin of canonical plant phytochromes. Nature Communications, 6, 1–12. 10.1038/ncomms8852

59. Li, W., and Morgan-Kiss, R. M. (2019). Influence of environmental drivers and potential interactions on the distribution of microbial communities from three permanently stratified Antarctic lakes. Frontiers in Microbiology, 10(MAY), 1–16. 10.3389/fmicb.2019.01067

60. Li, X., Huff, J., Crunkleton, D. W., and Johannes, T. W. (2023). Light intensity and spectral quality modulation for improved growth kinetics and biochemical composition of Chlamydomonas reinhardtii. Journal of Biotechnology, 375, 28–39. 10.1016/j.jbiotec.2023.08.007

61. Lizotte, M. P., and Priscu, J. C. (1992). Photosynthesis-irradiance relationships in phytoplankton from the physically stable water column of a perennially ice-covered Lake (Lake Bonney, Antarctica). Journal of Phycology, 28(2), 133–264.

62. Lu, D., Zhang, Y., Zhang, A., and Lu, C. (2022). Non-Photochemical Quenching: From Light Perception to Photoprotective Gene Expression. International Journal of Molecular Sciences, 23(2). 10.3390/ijms23020687

63. Lu, Y., and Xu, J. (2015). Phytohormones in microalgae: A new opportunity for microalgal biotechnology? Trends in Plant Science, 20(5), 273–282. 10.1016/j.tplants.2015.01.006

64. Marcelino, V. R., Cremen, M. C. M., Jackson, C. J., Larkum, A. A. W., and Verbruggen, H. (2016). Evolutionary dynamics of chloroplast genomes in low light: A case study of the endolithic green alga ostreobium quekettii. Genome Biology and Evolution, 8(9), 2939–2951. 10.1093/gbe/evw206

65. Masuda, T., and Fujita, Y. (2008). Regulation and evolution of chlorophyll metabolism. Photochemical and Photobiological Sciences, 7(10), 1131–1149. 10.1039/b807210h

66. Masuda, T., Tanaka, A., and Melis, A. (2003). Chlorophyll antenna size adjustments by irradiance in Dunaliella salina involve coordinate regulation of chlorophyll a oxygenase (CAO) and Lhcb gene expression. Plant Molecular Biology, 51(5), 757–771. 10.1023/A:1022545118212

67. McMinn, A., and Hegseth, E. N. (2004). Quantum yield and photosynthetic parameters of marine microalgae from the southern Arctic Ocean, Svalbard. Journal of the Marine Biological Association of the United Kingdom, 84(5), 865–871. 10.1017/S0025315404010112h

68. Melis, A., Murakami, A., Nemson, J. A., Aizawa, K., Ohki, K., and Fujita, Y. (1996). Chromatic regulation in Chlamydomonas reinhardtii alters photosystem stoichiometry and improves the quantum efficiency of photosynthesis. Photosynthesis Research, 47(3), 253–265. 10.1007/BF02184286

69. Merchant, S. S., Witman, G. B., Terry, A., Salamov, A., Fritz-Laylin, L. K., Marechal-Drouard, L., … Karpowicz, S. J. (2007). The Chlamydomonas Genome Reveals the Evolution of Key Animal and Plant Functions. Science, 318(5848), 245–250. Retrieved from www.sciencemag.org

70. Morgan, R. M., Ivanov, A. G., Priscu, J. C., Maxwell, D. P., and Huner, N. P. A. (1998). Structure and composition of the photochemical apparatus of the Antarctic green alga, Chlamydomonas subcaudata. Photosynthesis Research, 56(3), 303–314. 10.1023/A:1006048519302

71. Morgan-Kiss, R. M., Ivanov, A. G., and Huner, N. P. A. (2002). The Antarctic psychrophile, Chlamydomonas subcaudata, is deficient in State I-State II transitions. Planta, 214(3), 435–445. 10.1007/s004250100635

72. Morgan-Kiss, R. M., Ivanov, A. G., Pocock, T., Król, M., Gudynaite-Savitch, L., and Hüner, N. P. A. (2005). The antarctic psychrophile, Chlamydomonas raudensis Ettl (UWO241) (Chlorophyceae, Chlorophyta), exhibits a limited capacity to photoacclimate to red light. Journal of Phycology, 41(4), 791–800. 10.1111/j.1529-8817.2005.04174.x

73. Nakagawara, E., Sakuraba, Y., Yamasato, A., Tanaka, R., and Tanaka, A. (2007). Clp protease controls chlorophyll b synthesis by regulating the level of chlorophyllide a oxygenase. Plant Journal, 49(5), 800–809. 10.1111/j.1365-313X.2006.02996.x

74. Neale, P. J., and Priscu, J. C. (1995). The photosynthetic apparatus of phytoplankton from a perennially ice-covered antarctic lake: Acclimation to an extreme shade environment. Plant and Cell Physiology, 36(2), 253–263. 10.1093/oxfordjournals.pcp.a078757

75. Nick, S., Meurer, J., Soll, J., and Ankele, E. (2013). Nucleus-encoded light-harvesting chlorophyll a/b proteins are imported normally into chlorophyll b-free chloroplasts of arabidopsis. Molecular Plant, 6(3), 860–871. 10.1093/mp/sss113

76. Obryk, M. K., Doran, P. T., and Priscu, J. C. (2019). Prediction of Ice-Free Conditions for a Perennially Ice-Covered Antarctic Lake. Journal of Geophysical Research: Earth Surface, 124(2), 686–694. 10.1029/2018JF004756

77. Palenik, B., Grimwood, J., Aerts, A., Rouzé, P., Salamov, A., Putnam, N., … Grigoriev, I. V. (2007). The tiny eukaryote Ostreococcus provides genomic insights into the paradox of plankton speciation. Proceedings of the National Academy of Sciences of the United States of America, 104(18), 7705– 7710. 10.1073/pnas.0611046104

78. Pattanayak, G. K., Biswal, A. K., Reddy, V. S., and Tripathy, B. C. (2005). Light-dependent regulation of chlorophyll b biosynthesis in chlorophyllide a oxygenase overexpressing tobacco plants. Biochemical and Biophysical Research Communications, 326(2), 466–471. 10.1016/j.bbrc.2004.11.049

79. Pinnola, A. (2019). The rise and fall of Light-Harvesting Complex Stress-Related proteins as photoprotection agents during evolution. Journal of Experimental Botany, 70(20), 5527–5535. 10.1093/jxb/erz317

80. Platt, T., Gallegos, C. L., and Harrison, W. G. (1980). Photoinhibition of photosynthesis in natural assemblages of marine phytoplankton. Journal of Marine Research, 38(4), 687–701.

81. Poirier, M., Osmers, P., Wilkins, K., Morgan-Kiss, R. M., and Cvetkovska, M. (2023). Aberrant light sensing and motility in the green alga Chlamydomonas priscuii from the ice-covered Antarctic Lake Bonney. Plant Signaling and Behavior, 18(1). 10.1080/15592324.2023.2184588

82. Polle, J. E. W., Barry, K., Cushman, J., Schmutz, J., Tran, D., Hathwaik, L. T., … Magnuson, J. (2017). Draft nuclear genome sequence of the halophilic and beta-carotene-accumulating green alga Dunaliella salina strain CCAP19/18. American Society for Microbiology, 5(43), 17–19.

83. Possmayer, M., Gupta, R. K., Szyszka-Mroz, B., Maxwell, D. P., Lachance, M. A., Hüner, N. P. A., and Smith, D. R. (2016). Resolving the phylogenetic relationship between Chlamydomonas sp. UWO 241 and Chlamydomonas raudensis sag 49.72 (Chlorophyceae) with nuclear and plastid DNA sequences. Journal of Phycology, 52(2), 305–310. 10.1111/jpy.12383

84. Prochnik, S. E., Umen, J., Nedelcu, A. M., Hallmann, A., Miller, S. M., Nishii, I., … Rokhsar, D. S. (2010). Genomic analysis of organismal complexity in the multicellular green alga volvox carteri. Science, 329(5988), 223–226. 10.1126/science.1188800

85. Qin, R., Li, Y., Zhang, L., and Liu, J. (2021). The effect of salinity shock on the growth and rapid light curve of dunaliella salina. Aquaculture Research, 52(6), 2569–2579. 10.1111/are.15105

86. Ralph, P. J., and Gademann, R. (2005). Rapid light curves: A powerful tool to assess photosynthetic activity. Aquatic Botany, 82(3), 222–237. 10.1016/j.aquabot.2005.02.006

87. Rappaport, H. B., and Oliverio, A. M. (2023). Extreme environments offer an unprecedented opportunity to understand microbial eukaryotic ecology, evolution, and genome biology. Nature Communications, 14(1). 10.1038/s41467-023-40657-4

88. Raymond, J. A., and Morgan-Kiss, R. (2017). Multiple ice-binding proteins of probable prokaryotic origin in an Antarctic lake alga, Chlamydomonas sp. ICE-MDV (Chlorophyceae). Journal of Phycology, 53(4), 848–854. 10.1111/jpy.12550

89. Reeves, S., McMinn, A., and Martin, A. (2011). The effect of prolonged darkness on the growth, recovery and survival of Antarctic sea ice diatoms. Polar Biology, 34(7), 1019–1032. 10.1007/s00300-011-0961-x

90. Reinbothe, C., Bartsch, S., Eggink, L. L., Hoober, J. K., Brusslan, J., Andrade-Paz, R., … Reinbothe, S. (2006). A role for chlorophyllide a oxygenase in the regulated import and stabilization of light-harvesting chlorophyll a/b proteins. Proceedings of the National Academy of Sciences of the United States of America, 103(12), 4777–4782. 10.1073/pnas.0511066103

91. Revell, L. J. (2024). phytools 2.0: an updated R ecosystem for phylogenetic comparative methods (and other things). PeerJ, 12. 10.7717/peerj.16505

92. Roth, M. S., Cokus, S. J., Gallaher, S. D., Walter, A., Lopez, D., Erickson, E., … Niyogi, K. K. (2017). Chromosome-level genome assembly and transcriptome of the green alga Chromochloris zofingiensis illuminates astaxanthin production. Proceedings of the National Academy of Sciences of the United States of America, 114(21), E4296–E4305. 10.1073/pnas.1619928114

93. Sakuraba, Y., Tanaka, R., Yamasato, A., and Tanaka, A. (2009). Determination of a chloroplast degron in the regulatory domain of chlorophyllide a oxygenase. Journal of Biological Chemistry, 284(52), 36689–36699. 10.1074/jbc.M109.008144

94. Salomé, P. A., and Merchant, S. S. (2019). A series of fortunate events: Introducing chlamydomonas as a reference organism. Plant Cell, 31(8), 1682–1707. 10.1105/tpc.18.00952

95. Sasso, S., Stibor, H., Mittag, M., and Grossman, A. R. (2018). The natural history of model organisms from molecular manipulation of domesticated chlamydomonas reinhardtii to survival in nature. ELife, 7, 1–14. 10.7554/eLife.39233

96. Schumacher, I., Menghini, D., Ovinnikov, S., Hauenstein, M., Fankhauser, N., Zipfel, C., … Aubry, S. (2022). Evolution of chlorophyll degradation is associated with plant transition to land. Plant Journal, 109(6), 1473–1488. 10.1111/tpj.15645

97. Sherwell, S., Kalra, I., Li, W., McKnight, D. M., Priscu, J. C., and Morgan-Kiss, R. M. (2022). Antarctic lake phytoplankton and bacteria from near-surface waters exhibit high sensitivity to climate-driven disturbance. Environmental Microbiology, 24(12), 6017–6032. 10.1111/1462-2920.16113

98. Shui, J., Saunders, E., Needleman, R., Nappi, M., Cooper, J., Hall, L., … Stowe-Evans, E. (2009). Light-dependent and light-independent protochlorophyllide oxidoreductases in the chromatically adapting cyanobacterium fremyella diplosiphon UTEX 481. Plant and Cell Physiology, 50(8), 1507– 1521. 10.1093/pcp/pcp095

99. Sievers, F., Wilm, A., Dineen, D., Gibson, T. J., Karplus, K., Li, W., … Higgins, D. G. (2011). Fast, scalable generation of high-quality protein multiple sequence alignments using Clustal Omega. Molecular Systems Biology, 7(539). 10.1038/msb.2011.75

100. Smith, D. R., Cvetkovska, M., Hüner, N. P. A., and Morgan-Kiss, R. (2019). Presence and absence of light-independent chlorophyll biosynthesis among Chlamydomonas green algae in an ice-covered Antarctic lake. Communicative and Integrative Biology, 12(1), 148–150. 10.1080/19420889.2019.1676611

101. Stahl-Rommel, S., Kalra, I., D’Silva, S., Hahn, M. M., Popson, D., Cvetkovska, M., and Morgan-Kiss, R. M. (2021). Cyclic electron flow (CEF) and ascorbate pathway activity provide constitutive photoprotection for the photopsychrophile, Chlamydomonas sp. UWO 241 (renamed Chlamydomonas priscuii). Photosynthesis Research, (0123456789). 10.1007/s11120-021-00877-5

102. Stoecker, D. K., and Lavrentyev, P. J. (2018). Mixotrophic plankton in the polar seas: A pan-Arctic review. Frontiers in Marine Science, 5(AUG). 10.3389/fmars.2018.00292

103. Stolárik, T., Hedtke, B., Šantrůček, J., Ilík, P., Grimm, B., and Pavlovič, A. (2017). Transcriptional and post-translational control of chlorophyll biosynthesis by dark-operative protochlorophyllide oxidoreductase in Norway spruce. Photosynthesis Research, 132(2), 165–179. 10.1007/s11120-017-0354-2

104. Suzuki, S., Yamaguchi, H., Nakajima, N., and Kawachi, M. (2018). Raphidocelis subcapitata (=Pseudokirchneriella subcapitata) provides an insight into genome evolution and environmental adaptations in the Sphaeropleales. Scientific Reports, 8(1), 1–13. 10.1038/s41598-018-26331-6

105. Szyszka, B., Ivanov, A. G., and Hüner, N. P. A. (2007). Psychrophily is associated with differential energy partitioning, photosystem stoichiometry and polypeptide phosphorylation in Chlamydomonas raudensis. Biochimica et Biophysica Acta - Bioenergetics, 1767(6), 789–800. 10.1016/j.bbabio.2006.12.001

106. Szyszka-Mroz, B., Cvetkovska, M., Ivanov, A. G., Smith, D. R., Possmayer, M., Maxwell, D. P., and Hüner, N. P. A. (2019). Cold-Adapted Protein Kinases and Thylakoid Remodeling Impact Energy Distribution in an Antarctic Psychrophile. Plant Physiology, 180(3), 1291–1309. 10.1104/pp.19.00411

107. Szyszka-Mroz, B., Ivanov, A. G., Trick, C. G., and Hüner, N. P. A. (2022). Palmelloid formation in the Antarctic psychrophile, Chlamydomonas priscuii, is photoprotective. Frontiers in Plant Science, 13(August), 1–16. 10.3389/fpls.2022.911035

108. Szyszka-Mroz, B., Pittock, P., Ivanov, A. G., Lajoie, G., and Hüner, N. P. A. (2015). The antarctic psychrophile Chlamydomonas sp. UWO 241 preferentially phosphorylates a photosystem I-cytochrome b6/f supercomplex. Plant Physiology, 169(1), 717–736. 10.1104/pp.15.00625

109. Tanaka, A., Ito, H., Tanaka, R., Tanaka, N. K., Yoshida, K., and Okada, K. (1998). Chlorophyll a oxygenase (CAO) is involved in chlorophyll b formation from chlorophyll a. Proceedings of the National Academy of Sciences of the United States of America, 95(21), 12719–12723. 10.1073/pnas.95.21.12719

110. Tanaka, A., and Tanaka, R. (2019). The biochemistry, physiology, and evolution of the chlorophyll cycle. In Advances in Botanical Research (1st ed., Vol. 90). 10.1016/bs.abr.2019.03.005

111. Tanaka, R., and Tanaka, A. (2007). Tetrapyrrole biosynthesis in higher plants. Annual Review of Plant Biology, 58, 321–346. 10.1146/annurev.arplant.57.032905.105448

112. Tanaka, R., and Tanaka, A. (2005). Effects of chlorophyllide a oxygenase overexpression on light acclimation in Arabidopsis thaliana. Photosynthesis Research, 85(3), 327–340. 10.1007/s11120-005-6807-z

113. Untergasser, A., Cutcutache, I., Koressaar, T., Ye, J., Faircloth, B. C., Remm, M., and Rozen, S. G. (2012). Primer3-new capabilities and interfaces. Nucleic Acids Research, 40(15), 1–12. 10.1093/nar/gks596

114. Vedalankar, P., and Tripathy, B. C. (2019). Evolution of light-independent protochlorophyllide oxidoreductase. Protoplasma, 256(2), 293–312. 10.1007/s00709-018-1317-y

115. Wicke, S., Schneeweiss, G. M., dePamphilis, C. W., Müller, K. F., and Quandt, D. (2011). The evolution of the plastid chromosome in land plants: Gene content, gene order, gene function. Plant Molecular Biology, 76(3–5), 273–297. 10.1007/s11103-011-9762-4

116. Worden, A. Z., Lee, J., Mock, T., Rouzé, P., Simmons, M. P., Aerts, A. L., … Poliakov, A. (2009). Green evolution and dynamic adaptations revealed by genomes of the marina picoeukaryotes Micromonas. Science, 375(APRIL).

117. Wu, Y., Liao, W., Dawuda, M. M., Hu, L., and Yu, J. (2019). 5-Aminolevulinic acid (ALA) biosynthetic and metabolic pathways and its role in higher plants: a review. Plant Growth Regulation, 87(2), 357–374. 10.1007/s10725-018-0463-8

118. Yamasato, A., Nagata, N., Tanaka, R., and Tanaka, A. (2005). The N-terminal domain of chlorophyllide a oxygenase confers protein instability in response to chlorophyll b accumulation in Arabidopsis. Plant Cell, 17(5), 1585–1597. 10.1105/tpc.105.031518

119. Yong, S., Chen, Q., Xu, F., Fu, H., Liang, G., and Guo, Q. (2024). Exploring the interplay between angiosperm chlorophyll metabolism and environmental factors. Planta, 260(1), 1–16. 10.1007/s00425-024-04437-8

120. Zhang, L., Yang, C., and Liu, C. (2023). Revealing the significance of chlorophyll b in the moss Physcomitrium patens by knocking out two functional chlorophyllide a oxygenase. Photosynthesis Research, 158(3), 171–180. 10.1007/s11120-023-01044-8

121. Zhang, X., Cvetkovska, M., Morgan-Kiss, R., Hüner, N. P. A., and Smith, D. R. (2021). Draft genome sequence of the Antarctic green alga Chlamydomonas sp. UWO241. IScience, 24(2). 10.1016/j.isci.2021.102084

122. Zhang, X., Zheng, Z., He, Y., Liu, L., Qu, C., and Miao, J. (2020). Molecular Cloning and Expression of a Cryptochrome Gene CiCRY-DASH1 from the Antarctic microalga Chlamydomonas sp. ICE-L. Molecular Biotechnology, 62(2), 91–103. 10.1007/s12033-019-00225-y

