## Supplementary Figures and Tables for "Light quality affects chlorophyll biosynthesis and photosynthetic performance in Antarctic Chlamydomonas"

***Photosynthesis Research* Supporting Information**

**Author names:** Mackenzie Poirier, Kassandra Fugard, Marina Cvetkovska*

***Corresponding author:** Marina Cvetkovska

Department of Biology, University of Ottawa, 30 Marie-Curie Pr., Ottawa, ON, Canada, K1N 6N5

**Table S1** Sequences of species-specific primers used to measure CAO, LPOR, and CHLN gene expression and conserved primers used to confirm presence or absence of DPOR subunits in this study

| **Species** | **Gene** | **Forward (5’-3’)** | **Reverse (5’-3’)** |
| --- | --- | --- | --- |
| *C. priscuii* | LPOR | AAGTCTGTCCGCAACCTCTG | GGAACTTCTCGGGGTCCTTG |
|  | CAO1 | CAAGCAACAAGGTGTCGCTC | GTCCATCTTGAGCGCTACGA |
|  | CAO2 | CGAGGAAAACCCAACCGAGA | ACTCTGACTCGGTGTGCATG |
| ICE-MDV | LPOR | AACACCGTCAGTATGGCCAG | CCAGAGTCGATATTGGCGCT |
|  | CHLN | GGGGGACGGTTTTGACTCTC | TGACTCGCTACTGTTGTTGGT |
|  | CAO1 | GTGCTGTTCCGGGATGAGAA | GACAGTGACACCCTTGCAGA |
|  | CAO2 | AGGAGAACGGATGTGGAGGA | ACGCTGCTCTTCAGATGCTT |
| Conserved | CHLN | TTTGCTGAACCTCGTTATGCT | CGTGGTGCCATGCCTTCTAA |
|  | CHLL | TGTTGAAGCTGGTGGACCAC | CCCCCACAAACAACATCACC |

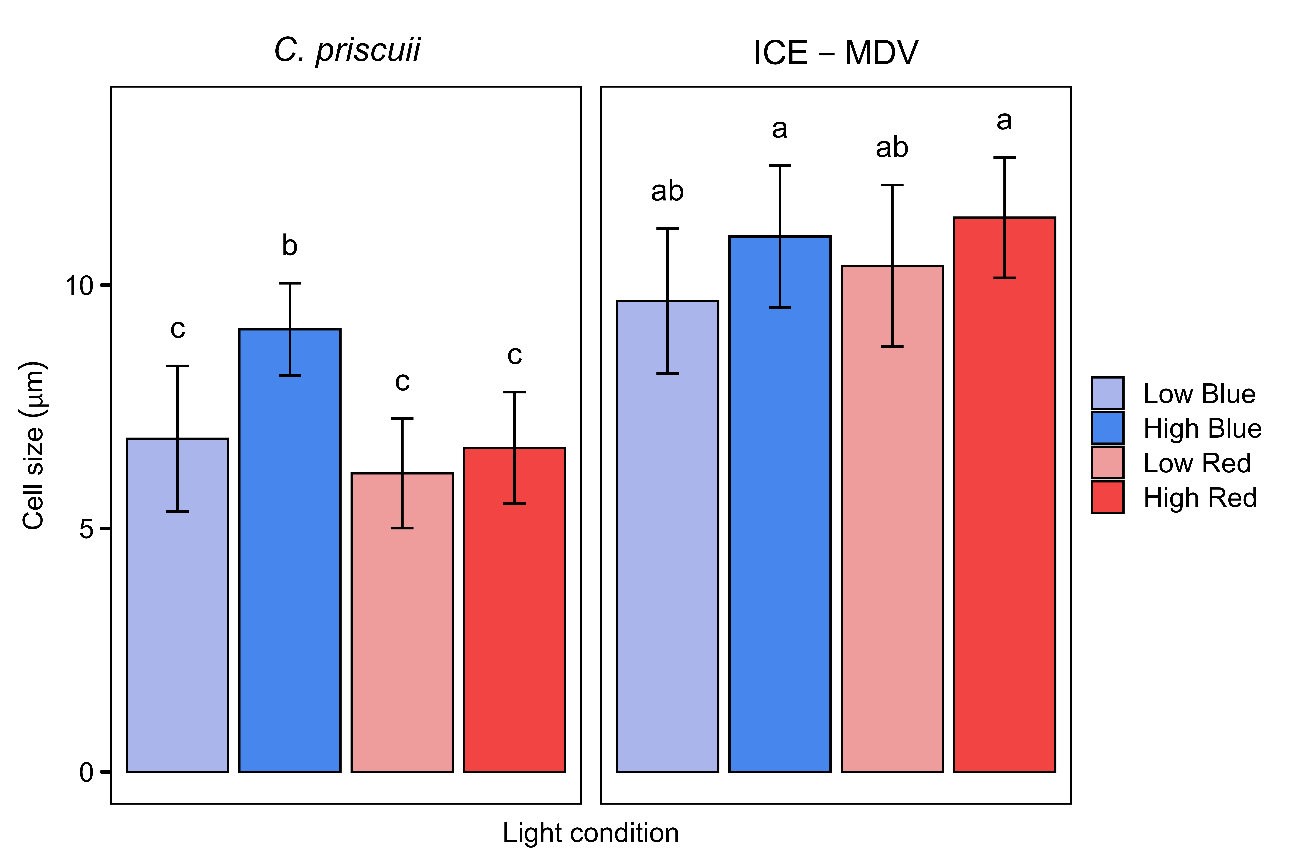

**Fig. S1** Average size of *C. priscuii* and ICE-MDV cells grown in different steady state light conditions. Statistically significant differences are represented as different letters as determined using Tukey’s post hoc test (p < 0.05). Values are means ± SD (n ≥ 9)

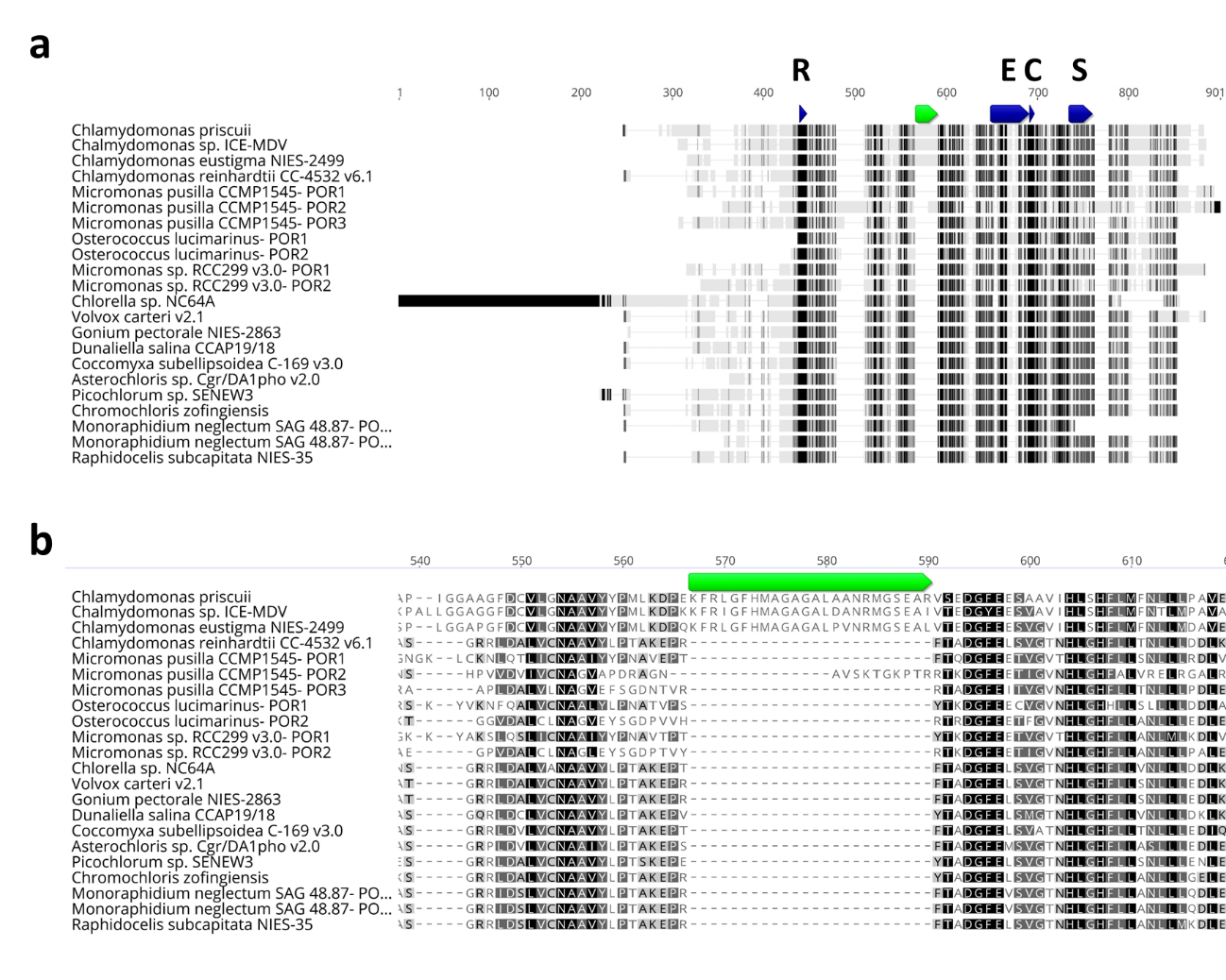

**Fig. S2** Alignment of LPOR sequences from representative chlorophyte species. Alignment of Chlorophyceae LPOR peptide sequences (a) demonstrates high conservation of catalytic domains involved in NADPH and Pchlide binding seen as dark blue boxes (Rossmann fold, R; Extended loop region, E; catalytic domain, C; substrate binding domain, S; Dong et al. 2020). A region of interest in the LPOR sequences present in the psychrophiles *C. priscuii* and ICE-MDV, as well as the acidophile *C. eustigma*, is highlighted in green. (b) A detailed view of the 24 amino acid insertion seen only in the algal extremophiles. In all cases, the degree of conservation between genes for each residue can be seen based on colour (black, fully conserved; dark gray, 80-100% similar; light gray, 60-80% similar; white, <60% similar). Sequences originate from the following species genomes: *Chlamydomonas priscuii* (Zhang et al. 2021)*, Chlamydomonas* sp. ICE-MDV (Raymond and Morgan-Kiss 2017), *Chlamydomonas eustigma* NIES-2499 (Hirooka et al. 2017), *Chlamydomonas reinhardtii* CC-4532 v6.1 (Craig et al. 2023), *Micromonas pusilla* CCMP1545 (Worden et al. 2009), *Osterococcus lucimarinus* (Palenik et al. 2007), *Micromonas* sp. RCC299 v3.0 (Worden et al. 2009), *Chlorella* sp. NC64A (Blanc et al. 2010), *Volvox carteri* v2.1 (Prochnik et al. 2010), *Gonium pectorale* NIES-2863 (Hanschen et al. 2016), *Dunaliella salina* CCAP19/18 (Polle et al. 2017)*, Coccomyxa subellipsoidea* C-169 v3.0 (Blanc et al. 2012), *Asterochloris* sp. Cgr/DA1pho v2.0 (Armaleo et al. 2019), *Pichlorum* sp. SENEW3 (da Roza et al. 2024), *Chromochloris zofingiensis* (Roth et al. 2017), *Monoraphidium neglectum* SAG 48.87 (Bogen et al. 2013), *Raphidocelis subcapitata* (Suzuki et al. 2018)

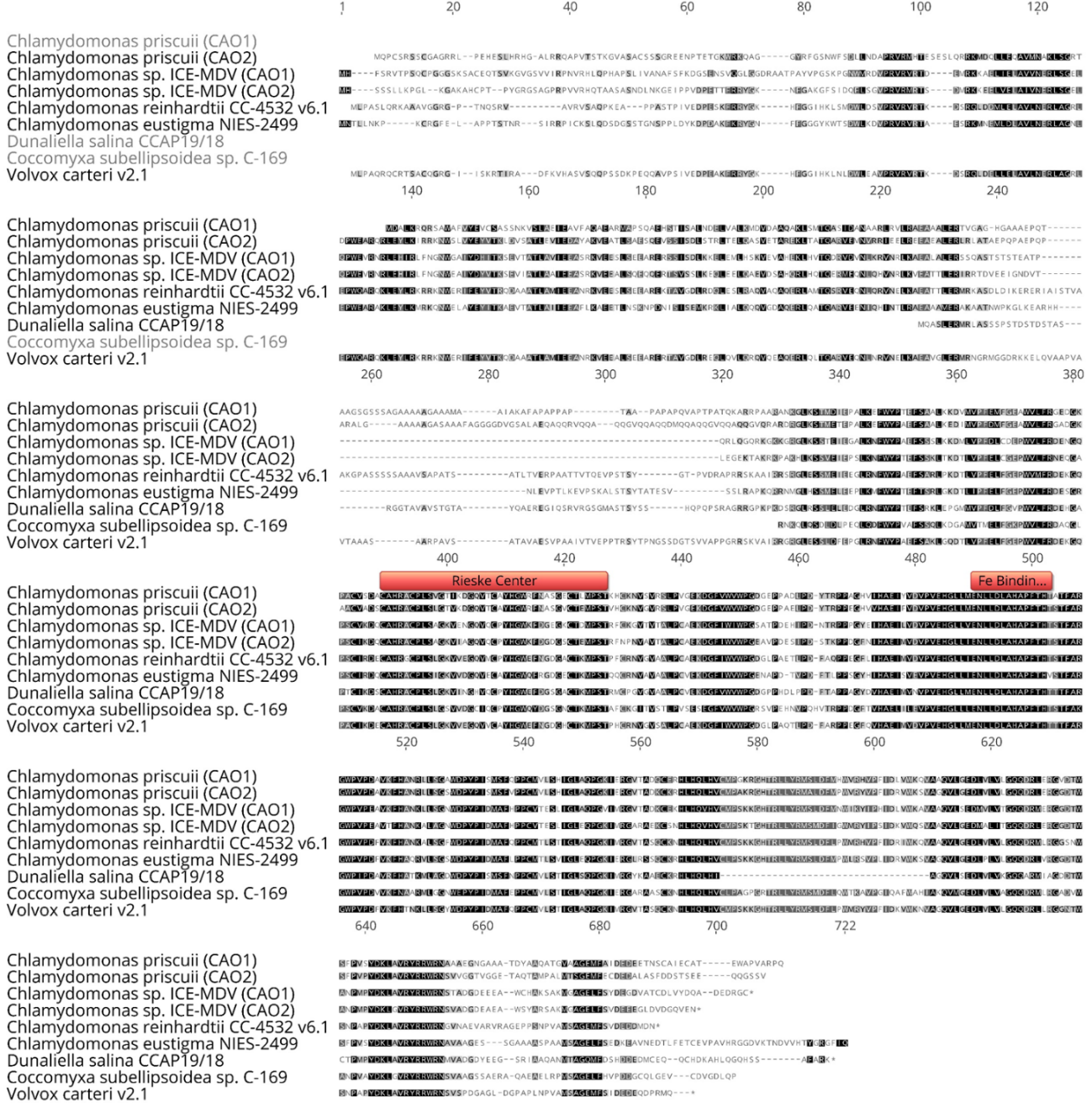

**Fig. S3** Multiple sequence alignment of CAO peptides. Predicted CAO peptide sequences identified by screening the genomes of *C. priscuii* and ICE-MDV were aligned with homologs from other chlorophyte species. Alignment done with the Geneious Global Alignment tool with free end gaps using the cost matrix Blosum62. Conserved Rieske and iron binding domains important for gene function (Tanaka et al. 1998) are highlighted in red. The degree of conservation between genes for each residue can be seen based on colour (black, fully conserved; dark gray, 80-100% similar; light gray, 60-80% similar; white, <60% similar). Sequences from the following species genomes: *Chlamydomonas priscuii* (Zhang et al. 2021)*, Chlamydomonas* sp. ICE-MDV (Raymond and Morgan-Kiss 2017), *Chlamydomonas reinhardtii* CC-4532 v6.1 (Craig et al. 2023), *Chlamydomonas eustigma* NIES-2499 (Hirooka et al. 2017), *Dunaliella salina* CCAP19/18 (Polle et al. 2017), *Coccomyxa subellipsoidea* C-169 v3.0 (Blanc et al. 2012), *Volvox carteri* v2.1 (Prochnik et al. 2010)

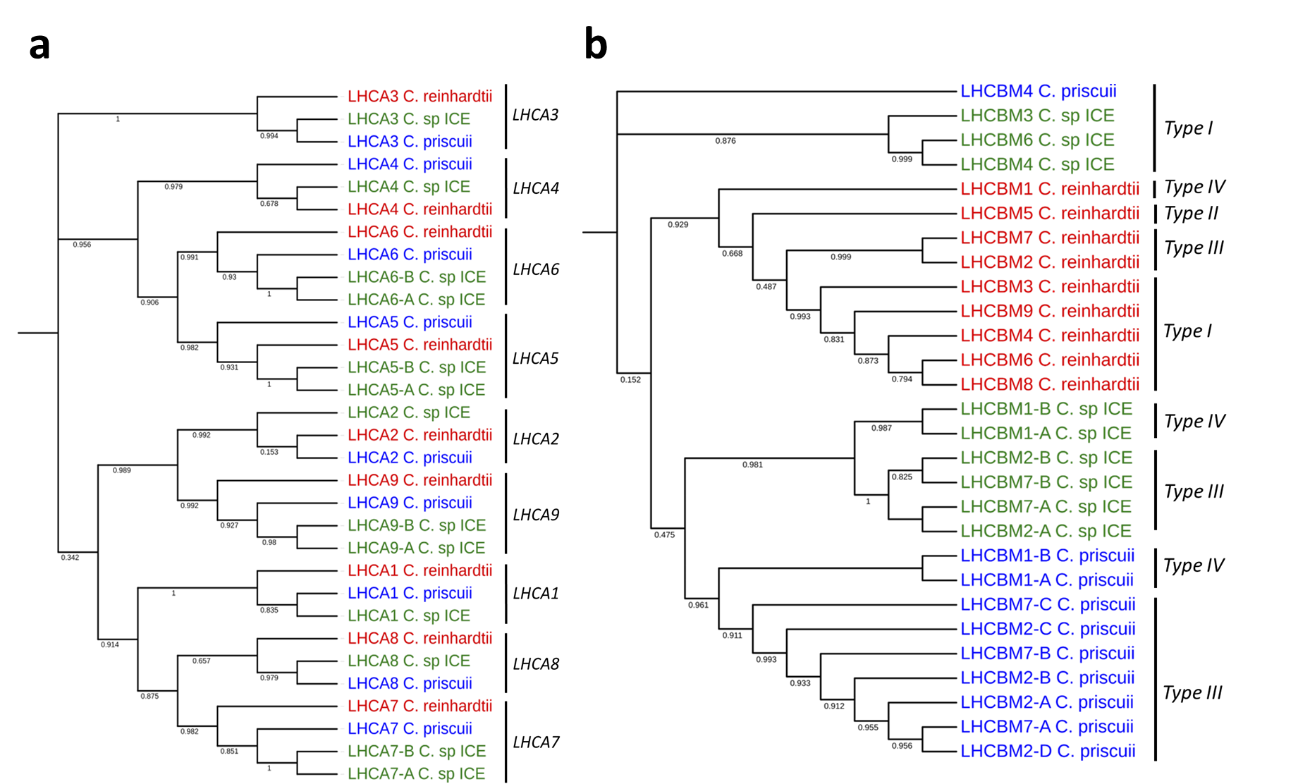

**Fig. S4** Phylogenetic analysis of LHCA (a) and LHCBM (b) genes from *C. priscuii* (blue), ICE-MDV (green) and *C. reinhardtii* (red), as determined by a maximum likelihood method with bootstrap values from 1000 iterations

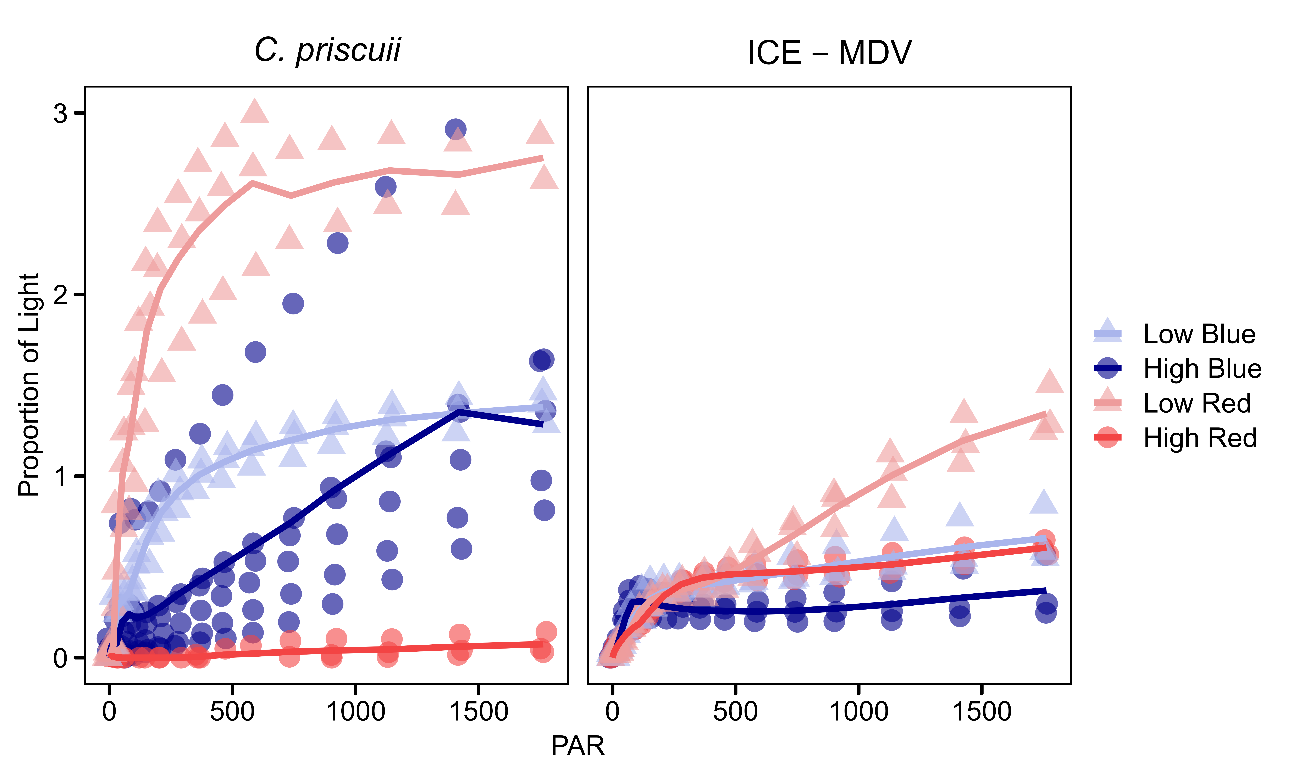

**Fig. S5** Nonphotochemical quenching (NPQ) at increasing light intensity measured during rapid light curve in *C. priscuii* and ICE-MDV cultures grown in different steady state light conditions. Points represent individual biological replicates and lines represent mean (n ≥ 3)

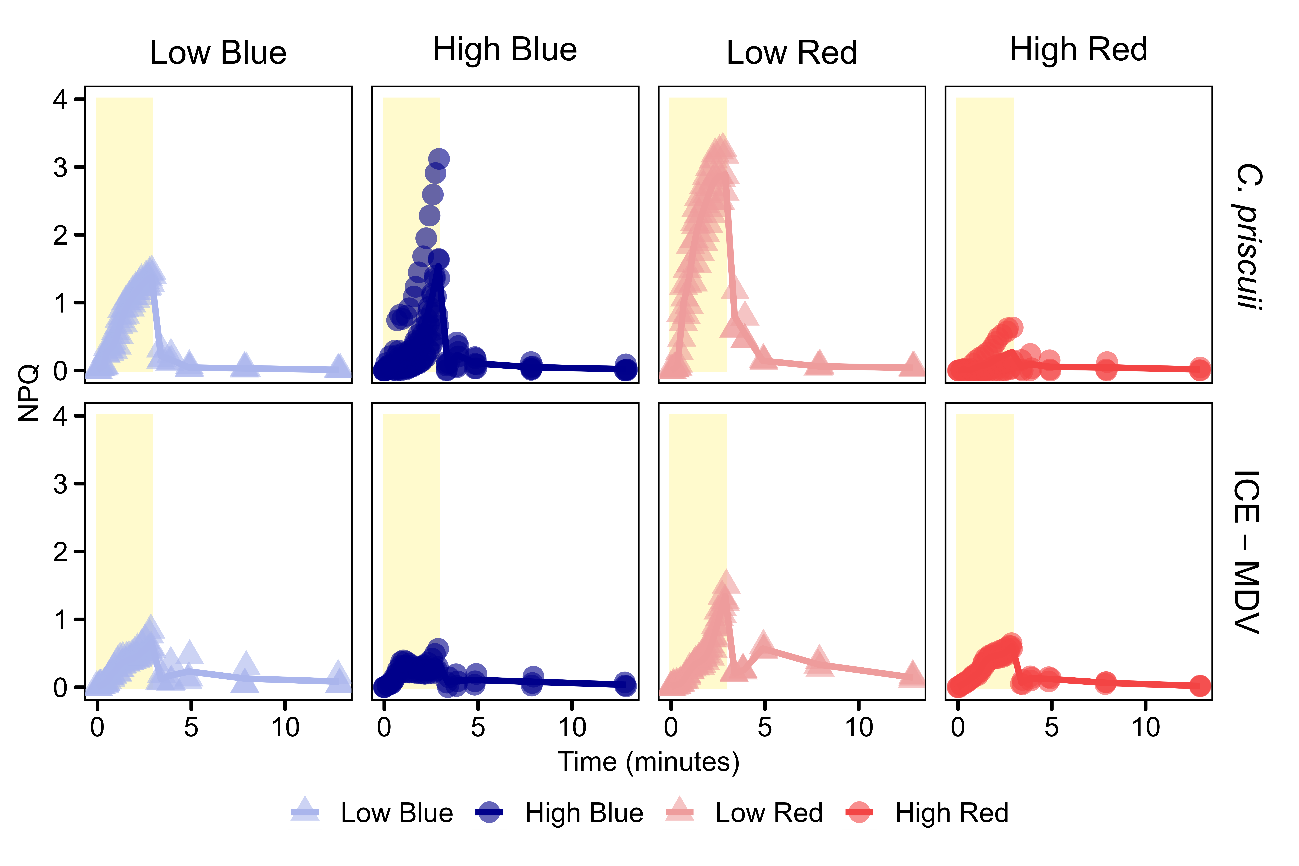

**Fig. S6** NPQ rates during rapid light curve and dark recovery effects over time in *C. priscuii* and ICE-MDV cultures grown in different steady state light conditions. The yellow box indicates the light time period during the rapid light curve which was followed by a 10 minute dark recovery period. Points represent individual biological replicates and lines represent mean (n ≥ 3)
